# Vacuolar pH regulates clathrin-mediated endocytosis through TORC1 signaling during yeast replicative aging

**DOI:** 10.1101/2024.11.28.625547

**Authors:** Kenneth Gabriel Antenor, Jaime Lee-Dadswell, Nina Grishchenko, Shaimaa Swaleh, Allie Spangaro, Mojca Mattiazzi Usaj

**Author notes:** **Address correspondence to:** Mojca Mattiazzi Usaj.

## Abstract

Clathrin-mediated endocytosis (CME) is a critical cellular process that regulates nutrient uptake, membrane composition and signalling. While cellular aging is associated with functional changes across many cellular components contributing to the collective decline in cellular function, little is known about how it affects CME. Here we show that CME dynamics are significantly altered during replicative aging in budding yeast, with older cells having slower assembly of early and coat CME modules, resulting in longer endocytic turnover and reduced cargo internalization. This change in CME dynamics is mother cell-specific and is not observed in daughter cells. We identified vacuolar pH, a key driver of aging phenotypes in budding yeast, as a central player in this modulation of CME dynamics during aging. Perturbing vacuolar pH in young cells mimics aging-like CME dynamics, while maintaining an acidic vacuolar pH in aging cells preserves CME dynamics typical of young cells. Finally, we demonstrate that the vacuolar pH effect on CME is regulated through TORC1 via the effector kinase Npr1. These findings establish vacuolar pH as a critical regulator of CME during cellular aging, and strengthen its role in the overall cellular aging process in budding yeast.

## INTRODUCTION

Clathrin-mediated endocytosis (CME), a fundamental bioprocess highly conserved among eukaryotes, is required for the internalization of cargo such as transporters, lipids, and receptors from the plasma membrane, and is hence crucial for homeostasis of broader cellular functions, including response to stress and adaptation to nutrient availability (Fiore & Zastrow, 2014). As such, it is not surprising that CME is implicated in diverse physiological processes, including antigen presentation, neuronal function and progression of diseases, such as neurodegeneration, hypercholesterolemia and cancer metastasis (Garcia et al., 2001; Mellman & Yarden, 2013; Tessneer et al., 2013).

The use of budding yeast, *Saccharomyces cerevisiae*, as a model organism has been instrumental in advancing our understanding of CME, providing key insights into the molecular machinery and regulation of this essential cellular process. In budding yeast, CME is tightly regulated and involves the sequential recruitment of distinct functional modules, including the early module, coat module, WASP/myosin, actin, and amphiphysin modules, which assemble at discrete plasma membrane sites known as CME patches (Lu et al., 2016). The temporal architecture of CME consists of two distinct phases: the variable phase, which is characterized by variable protein lifetimes, and the regular phase (Pedersen et al., 2020). The transition from variable to regular phase is defined as “cargo-sensitive”, thereby positioning cargo (including plasma membrane amino acid transporters) as an essential component for CME progression (Carroll et al., 2012; Pedersen et al., 2020; Tolsma et al., 2020). CME sites are initiated by the early module proteins (Maldonado-Báez et al., 2008; Peng et al., 2015; Stimpson et al., 2009), followed by coat proteins that coordinate F-actin assembly through recruitment of actin nucleation factors (Barker et al., 2007; Feliciano & Pietro, 2012; Sun et al., 2015; Tolsma et al., 2018). The polymerized F-actin, together with myosin and other actin module proteins, generates the pulling force necessary to bend the plasma membrane inward (Idrissi et al., 2002; Okreglak & Drubin, 2007; Pedersen & Drubin, 2019; Sun et al., 2006). Finally, amphiphysins are recruited to the vesicle neck, completing the formation of the cargo-containing vesicle (Balguerie et al., 1999; Lombardi & Riezman, 2001; Myers et al., 2016; Palmer et al., 2015; Youn et al., 2010). Despite the critical role CME plays in maintaining cellular homeostasis, the effects of cellular aging on CME, specifically how aging may impact the regulation and spatio-temporal dynamics of CME, remain largely unknown.

Research on replicative aging in budding yeast has been instrumental in elucidating the hallmarks of eukaryotic aging (Janssens & Veenhoff, 2016). Replicative aging in budding yeast is studied by analyzing a cell’s limited replicative lifespan, defined as the number of daughter cells a mother cell produces before its death (Longo et al., 2012; Mortimer & Johnston, 1959). In wildtype cells, the average replicative lifespan is around 25-30 divisions (Liu et al., 2015; McCormick et al., 2015). During mitosis in budding yeast, the mother cell produces a rejuvenated daughter cell through asymmetric partitioning of cellular components, with diffusion barriers forming around the bud neck to help establish this mother-daughter asymmetry (Clay et al., 2014; Erjavec & Nyström, 2007; Higuchi-Sanabria et al., 2014; Shcheprova et al., 2008). Consequently, the newly-produced daughter cells have full replicative potential, unlike the mother cells, which exhibit reduced replicative potential and cellular functioning. Budding yeast’s replicative aging is thus an effective model to study cellular aging of mitotic cells in higher eukaryotes.

Cellular homeostasis also depends on proper functioning of organelles and other cellular components. During aging, many organelles undergo structural and functional changes contributing to the decline in overall cellular function. Loss of organelle integrity is thus considered one of the drivers of cellular aging and senescence, and is linked to several age-related pathologies, including neurodegeneration, bone deterioration, and reduced fertility (Babayev & Duncan, 2022; Hou et al., 2019; Qu et al., 2024). Several studies have shown that aging yeast mother cells have altered function of various organelles, particularly vacuoles (Henderson et al., 2014; Hughes & Gottschling, 2012; Hughes et al., 2020). Vacuoles, the yeast equivalent of mammalian lysosomes, are involved in a wide array of cellular processes, such as endocytosis, autophagy, and metabolite storage (Li & Kane, 2009). Their luminal pH is slightly acidic compared to the cytosol, and is maintained by the multimeric vacuolar ATPase (V-ATPase) which translocates protons into the vacuolar lumen (Forgac, 1999; Graham & Stevens, 1999; Liu & Kane, 1996). Acting as the endpoint for endocytic trafficking, vacuoles process cargo from the plasma membrane through the numerous vacuolar enzymes. Vacuoles also sequester metabolites, such as heavy metals like zinc and iron, as well as amino acids such as cysteine, to prevent their potentially harmful accumulation in the cytosol (MacDiarmid et al., 2000; Miseta et al., 1999; Russnak et al., 2001).

Vacuoles exhibit distinct changes throughout replicative aging. Older budding yeast cells tend to have multilobed vacuoles (Mattiazzi Usaj et al., 2020). In addition, during replicative aging, the vacuolar pH of aging mother cells becomes less acidic by the 4^th^ division in contrast to the daughter cell with an intact vacuolar pH (Hughes & Gottschling, 2012). This age-associated difference in vacuolar pH has been attributed to the asymmetric segregation of the plasma membrane proton antiporter, Pma1, a P2-type H^+^-ATPase (Serrano, 1978; Serrano et al., 1986). During replicative aging, Pma1 preferentially accumulates at the plasma membrane of mother cells leading to a less acidic cytosolic pH, and in turn a less acidic vacuolar pH, compared to their daughter cells (Henderson et al., 2014). Several studies also indicate that vacuolar pH dysfunction precedes mitochondrial defects during aging (Chen et al., 2020; Hughes & Gottschling, 2012; Hughes et al., 2020). During yeast replicative aging, alkalinized vacuoles lose their ability to properly import amino acids, particularly cysteine. This leads to mitochondrial damage as excess cytosolic cysteine undergoes oxidation, producing ROS that in turn deplete iron-sulfur clusters required for mitochondrial respiration (Chen et al., 2020; Hughes et al., 2020).

Vacuoles also participate in several signaling pathways, and vacuole-based signaling plays a crucial role in modulating the dynamics of CME. Specifically, the highly conserved Target of Rapamycin 1 (TORC1) pathway, which plays a crucial role in nutrient sensing and regulation of cellular growth (Aronova et al., 2007; Jorgensen et al., 2004), regulates CME through Npr1, a kinase that inhibits the arrestin-like trafficking adaptors known as alpha arrestins, which are required for recognition, ubiquitylation and internalization of amino acid transporters via CME (Craene et al., 2001; Lin et al., 2008; MacGurn et al., 2011; Schmidt et al., 1998). However, if and how vacuolar pH and TORC1 signalling affect CME during aging remains unexplored.

In this study, we investigated how CME dynamics are altered during cellular aging in budding yeast. We found that aging cells exhibit slower CME vesicle formation and delayed recruitment of later-arriving functional modules, leading to reduced cargo internalization. We identified vacuolar pH as a key regulator of these age-related defects: young cells with alkalinized vacuoles display CME lifetimes similar to those of older cells; on the other hand, restoration of vacuolar acidity through caloric restriction or deletion of a V-ATPase disassembly factor led to normal, young cell-like CME dynamics in older cells. We also show that introduction of a hyperactive form of TORC1, deletion of Npr1, or overexpression of alpha-arrestins all result in similar CME dynamics between young and older cells. We propose that downregulation of the vacuole-based TORC1 signaling complex negatively regulates CME dynamics under conditions of compromised vacuolar pH, such as during replicative aging.

## MATERIALS AND METHODS

### Yeast strains

*S. cerevisiae* strains used in this study are listed in Supplementary Table 1.

### Strain Construction and Growth

Yeast strains were constructed using two methods: i) transformation of either high-copy 2µ plasmid (Molecular barcoded yeast open reading frame (MoBY-ORF) library 2.0) (Magtanong et al., 2011), or polymerase chain reaction (PCR) product containing yomNeonGreen, yomScarlet-I or mCherry flanked by ∼45 bp of homology to the target open reading frame (ORF), using the lithium acetate protocol (Gietz & Schiestl, 2007); or by ii) genetic crossing and selection of haploid meiotic progeny bearing the genotype of interest. Construction method is indicated in Table 1 with * (homologous recombination of PCR product) and ** (genetic crossing). For all constructed strains, fluorescent-protein-tagged ORFs were verified by genotype PCR using oligonucleotides with homology regions up- and downstream of the tagged site along with fluorescence microscopy. Plasmids and oligonucleotides used in the study are listed in Supplementary Table 1.

All strains were grown in synthetic complete medium (SC) (0.67% (w/v) YNB with (NH_4_)_2_SO_4_, complete amino acid supplement (2 g/L), 2% (w/v) glucose) unless otherwise indicated. Strains carrying 2µ vectors for overexpression of alpha arrestins were grown in synthetic dextrose leucine drop-out medium (SD-Leu) (0.67% (w/v) YNB with (NH_4_)_2_SO_4_, amino acid supplement lacking leucine (DO-Leu; 2 g/L), 2% (w/v) glucose) supplemented with the antibiotic geneticin (200 mg/mL; Bioshop).

### Bud scar staining and determination of replicative age

To determine the replicative age of individual cells, an equivalent to OD_600_=1 of mid-log phase culture grown in SC medium was harvested, washed with phosphate buffered saline (PBS; Bioshop), and resuspended in CF640R wheat germ agglutinin (WGA) conjugate (CF640R WGA; Biotium) at a final concentration of 0.5 μg/mL in PBS. Cultures were incubated with constant agitation for 20 minutes in the dark at room temperature. Following incubation, the cells were washed three times with PBS and resuspended in the appropriate imaging medium. WGA-stained cells were immobilized to Concanavalin A- (ConA, 1mg/mL; BioLynx) coated coverslips and sealed to glass slides with vacuum grease (Dow Corning) for experiments related to Fig. 1. For the rest of the experiments, WGA-stained cells were adhered to ConA-coated 384-well imaging microplates (CellCarrier Ultra, PerkinElmer). The replicative age of individual cells was determined by binning based on WGA intensity percentiles for experiments related to Figs. 1 and S1A-E, and manually counting visible bud scars from maximum intensity z-projection images for the rest of the experiments.

**Fig. 1.**
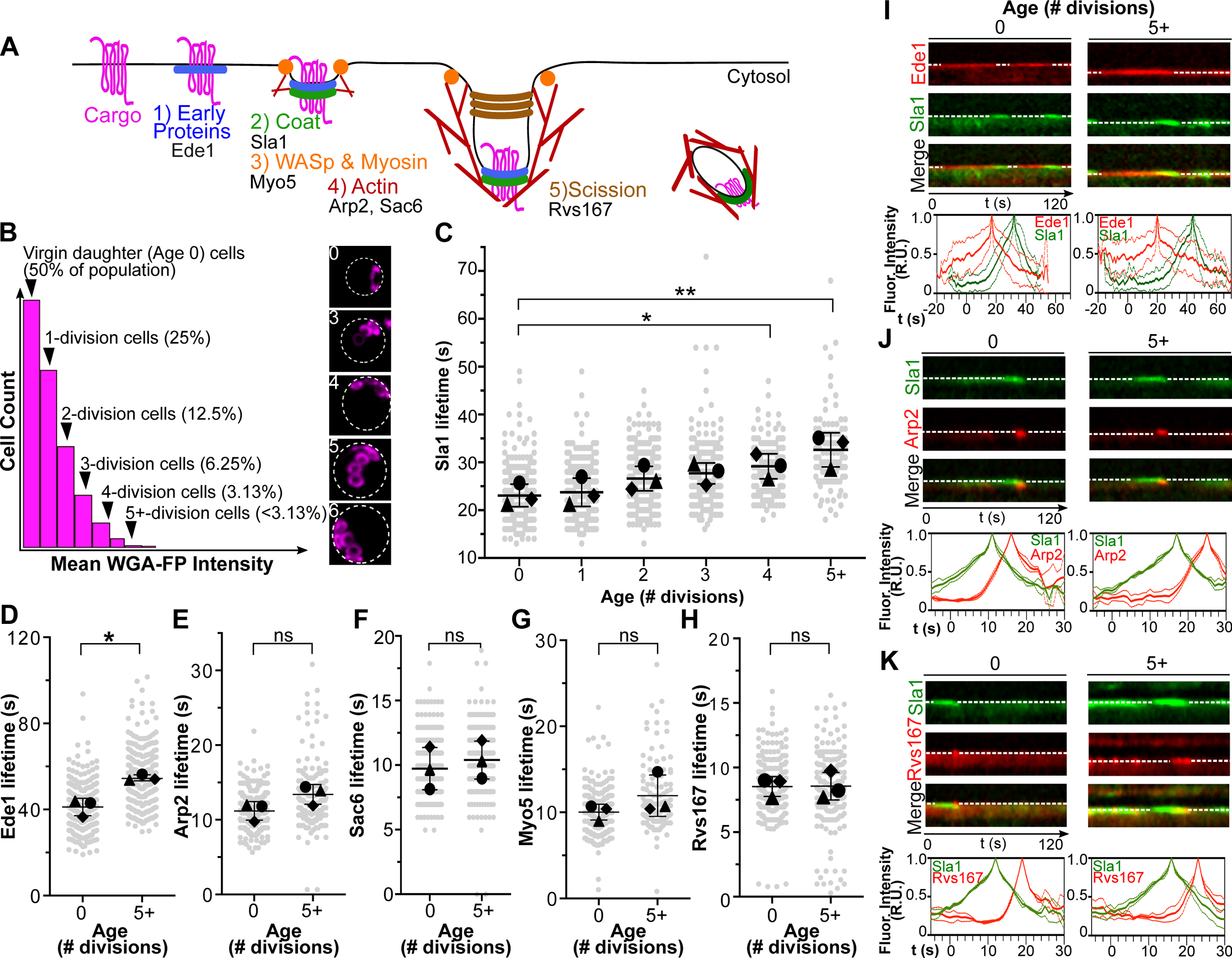
Lifetimes of proteins from early and coat module are prolonged in older cells. A) Schematic diagram of clathrin-mediated endocytosis in budding yeast. Functional modules are shown along with representative proteins analyzed in this study. B) Geometric distribution of replicative age in an asynchronous cell population. Briefly, age 0 cells represent 50% of the population, age 1 cells the next 25% percentile, age 2 cells the following 12.5% percentile, and so on. Example micrographs of cells with increasing numbers of WGA-fluorophore-stained bud scars are also shown. WGA intensity is directly proportional to replicative age. C) Patch lifetime of Sla1 in cells of increasing replicative age; and D-H) Patch lifetimes of Ede1, Arp2, Sac6, Myo5 and Rvs167 in cells of replicative age 0 and 5+. Replicative age of individual WGA-stained cells was assigned using the WGA intensity distribution within a population: top 50% representing age 0 cells, next 25% representing age 1 cells, and so on with bottom 3.13% consisting of age 5+ cells. For all experiments, patch lifetime data are shown as mean values ± standard deviation from three biological replicates (circle = replicate 1; triangle = replicate 2; diamond = replicate 3), overlayed on a swarm plot of individual patch measurements. At least 100 patches were quantified from 10-20 cells per strain for each age group and for each replicate. Statistical significance was assessed using Welch’s t-tests. *, p<0.05; **, p<0.01; ns, not-significant. I-K) Representative kymographs of Sla1 with Ede1 (I), Arp2 (J) and Rvs167 (K) from cells of replicative age 0 and 5+. Kymographs are shown with a dashed line representing the plasma membrane with the cytosol at the bottom. Fluorescence intensity quantification of Sla1 with Ede1 and representative late CME proteins Arp2 and Rvs167 as a function of time are also shown below each kymograph. Plots are shown as a statistical summary of mean (solid line) ± standard deviation (dotted line) from three biological replicates each with n>50 patches analyzed for each age category.

### Raffinose-induced transporter endocytosis

To determine the effect of replicative aging on cargo internalization, strains expressing Mup1-GFP and Sla1-tdTomato were kept in synthetic dextrose methionine drop-out medium (SD-Met) (0.67% (w/v) YNB with (NH_4_)_2_SO_4_, amino acid supplement lacking methionine (DO-Met; 2 g/L), 2% (w/v) glucose) to sequester Mup1-GFP on the plasma membrane.

For raffinose-induced Mup1-GFP endocytosis, overnight cultures were prepared in SD-Met and then grown to mid-log phase in the same medium (Laidlaw et al., 2021). Bud scar staining of the mid-log culture was conducted as described above. WGA-stained cells were resuspended with synthetic raffinose methionine drop-out medium (SRaff-Met + 2% (w/v) raffinose) and transferred to a ConA-treated 384-well imaging microplate. Z-stacks were acquired within 40 minutes of the media change as described below.

### V-ATPase inhibition and vacuole staining

To determine the effect of V-ATPase inhibition on CME dynamics, strains expressing Sla1-mCherry were grown to mid-log phase in SC medium, washed once with PBS, and resuspended in SC medium containing Concanamycin A (Sigma) at the indicated final concentrations. After 15 min incubation at 30°C, cells were washed three times with PBS and stained with quinacrine to visualize vacuole acidity as previously described (Morano & Klionsky, 1994). Briefly, cells were washed once with the uptake buffer (YPAD (1% (w/v) yeast extract, 2% (w/v) peptone, 0.012% (w/v) adenine sulfate, 2% (w/v) glucose) + 100 mM HEPES, pH 7.6), resuspended in 100 µL of uptake buffer containing 200 µM quinacrine (Sigma) and incubated at 30°C for 10 minutes, followed by a 5 min incubation on ice. Cells were washed three times with an ice-cold wash buffer (100 mM HEPES + 2% glucose, pH 7.6) followed by WGA staining as described above. Quinacrine- and WGA-stained cells were transferred to either ConA-coated 384-well microplates or slides, and imaged in SC medium.

### CME inhibition by Latrunculin A

To determine if CME inhibition results in a change in vacuole acidity in young cells, a strain expressing Sac6-yomScarlet-I was grown to mid-log phase in SC medium, washed once with PBS, and stained with quinacrine and WGA as described above. Stained cells were resuspended in either control or SC medium containing 25µM Latrunculin A (LatA) (Abcam), and imaged within 40 minutes of resuspension in LatA-containing medium.

### Caloric Restriction

To determine whether caloric restriction can restore CME dynamics during replicative aging, strains expressing Sla1-yomNeonGreen were grown overnight, and then grown to mid-log phase in SC medium containing either 2% (control) or 0.5% (caloric restriction) glucose. An equivalent of OD_600_=1.0 of mid-log culture was harvested, washed once with PBS, resuspended in the growth medium and incubated for 2 hours at 30°C before staining with WGA as described above. WGA-stained cells were resuspended in the same growth medium (control or caloric restriction) for imaging.

### TORC1 inhibition by rapamycin

To determine whether TORC1 regulates CME dynamics during replicative aging, strains expressing Sla1-yomNeonGreen were grown to mid-log phase in SC medium at 30°C with shaking, washed once with PBS, and resuspended in SC medium containing 200 ng/mL rapamycin (Bioshop). After incubation for 2 hours at 30°C to inhibit TORC1 activity, cells were washed three times with PBS, stained with WGA as described above, resuspended in fresh SC medium containing 200 ng/mL rapamycin, and imaged immediately.

### Live-cell Fluorescence microscopy

For experiments for Fig. 1 and S1A-E, acquisition of z-stacks and two-colour time-lapse movies was done at room temperature, unless otherwise indicated, on a Leica DMi8 Microscope connected to an Andor iXON 897 EMCCD camera, using a 1.4 NA 63x oil-immersion objective. Z-stacks of 17 slices with 0.25 μm step size were acquired using uniform 50% laser power. The acquisition settings (excitation wavelength / exposure time) for the fluorophores were: 668 nm/100 ms for CF640R WGA; 455 nm/100 ms for Hta2-TagBFP; 519 nm/400 ms for yomNeonGreen; and 617 nm/400ms for yomScarlet-I. Two-colour time-lapse movies of yomNeonGreen and yomScarlet-I were acquired for a duration of 2 min with a frame rate of 1 image per second using 100 ms exposure and 50% laser power.

For the experiments associated with Figs. 2-6, S1F-K, and S2-S5, WGA-stained cells were adhered to ConA-coated 384-well imaging microplates (CellCarrier Ultra, PerkinElmer). Opera Phenix automated spinning disc confocal microscope (PerkinElmer) equipped with Harmony High-Content Imaging and Analysis software was used to acquire both z-stacks and single-channel live-cell movies at room temperature, unless otherwise indicated. Four-channel z-stacks of 5 slices with 1.0 μm spacing were acquired using a 60X water-immersion objective and a 405/488/567/640 nm primary dichroic mirror. yomNeonGreen/GFP was excited using 488 nm laser (exposure, 800 ms; laser power, 100%); yomScarlet/mCherry was excited using 561 nm laser (800 ms, 50%); TagBFP was excited using 405 nm laser (800 ms, 50%); and, CF640R WGA was excited using 640 nm laser (800 ms, 50%). Time-lapse movies were acquired for a duration of 2 min with a frame rate of 1 image per second using 900 ms exposure and 15% laser power for yomNeongreen-, yomScarlet- and mCherry-tagged proteins.

**Fig. 2.**
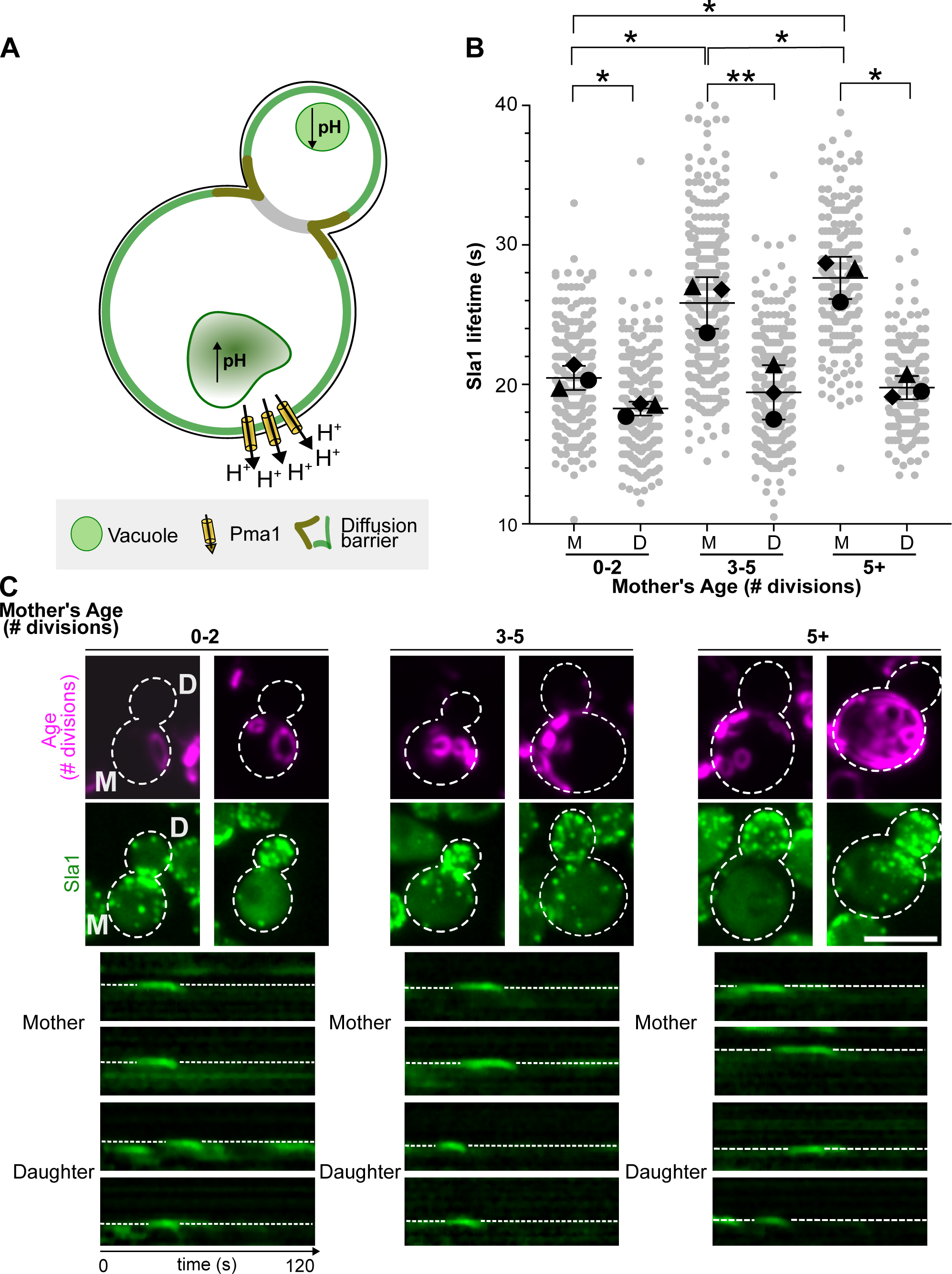
Daughter cells do not display prolonged Sla1 patch lifetimes. A) Asymmetric segregation during yeast replicative aging. During cell division, mother cells retain aging factors, including alkalinized vacuoles and Pma1, due to the diffusion barriers at the bud neck. Daughter cells, however, have a functional acidic vacuole, lack aging factors, and are rejuvenated with full replicative potential. B) Quantification of Sla1 patch lifetimes in mother and daughter cells. Sla1 patch lifetimes were measured in mother cells (*M*) of different replicative ages and their immediate buds/daughters (*D*). Mother cells were grouped into three replicative age categories (0-2, 3-5, and 5+) based on WGA-stained bud scar counts. Data is reported as the statistical summary of the mean patch lifetime values ± standard deviation from three biological replicates. Means of individual biological replicates are represented with a circle, triangle, and diamond). Individual patch measurements are displayed as a swarm plot. At least 50 patches were analyzed per replicate and age group. Statistical significance was assessed using paired t-tests. *, p<0.05; **, p<0.01; ns, not-significant. C) Representative micrographs and kymographs of Sla1 patches. Top panel: Micrographs showing actively budding mother cells of different replicative ages. Bottom panel: Kymographs of individual Sla1 patches obtained from mother and daughter cells. Plasma membrane is indicated by dashed lines, with the cytosol at the bottom. Scale bar: 5 µm

### Image and Data Analysis

For CME patch lifetime measurements (all Figures except Fig.3, S2 and 7), kymographs were generated using the Radial Reslice plugin in ImageJ (Schneider et al., 2012). A horizontal line (line width = 1) was drawn from the start to the end of a patch’s kymograph to determine the lifetime. Patches that successfully internalized (i.e., formed a cytosolic-generated vesicle) were visually identified with an end region that invaginates toward the cell interior. The duration of Sla1-yomNeonGreen before the arrival of ORF-yomScarlet-I (Fig. S1B-E) was measured by subtracting ORF-yomScarlet-I lifetime from Sla1-yomNeonGreen’s lifetime. For the analysis of Ede1-yomNeongreen duration before Sla1-yomScarlet-I arrival (Fig. S1A), the same calculation was conducted using Ede1 lifetime as reference. At least 100 patches were analyzed for each age category in each of the three biological replicates.

**Fig. 3.**
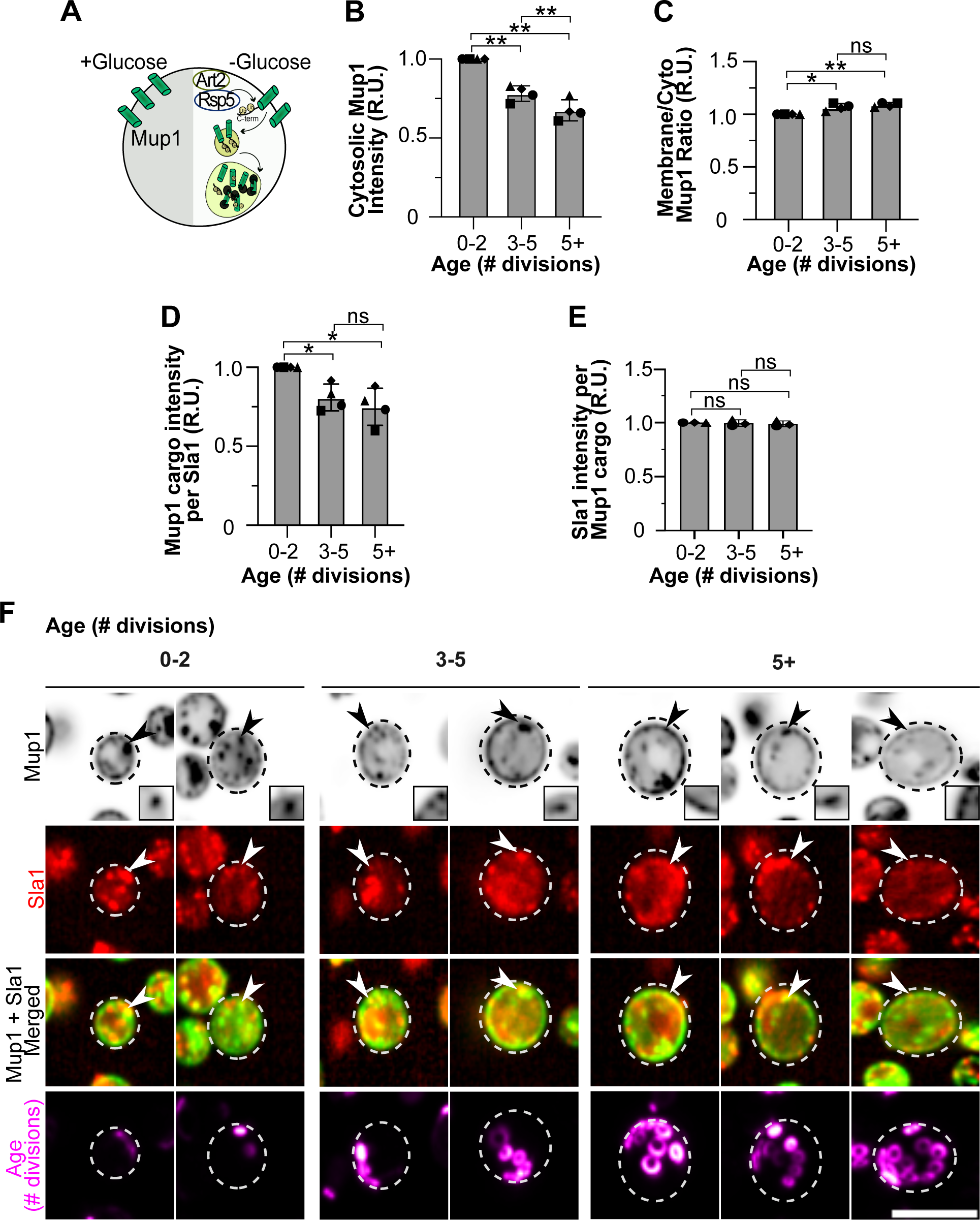
Older cells internalize less endocytic cargo compared to young cells. A) Schematic representation of glucose starvation-induced internalization of PM transporters. Under normal glucose conditions, PM transporters such as the methionine permease, Mup1, are sequestered at the cell surface. During glucose depletion/starvation conditions, Mup1 is ubiquitinated by the Art2-Rsp5 alpha-arrestin-ubiquitin ligase complex leading to its internalization via CME. Internalized Mup1 is then trafficked to the endosomal system and degraded in the vacuole. B - E) Quantitative analysis of Mup1 internalization in cells from varying replicative age groups. (B) Cytosolic, (C) Ratio of Mup1-GFP at membrane vs cytosol, (D) Mup1-GFP in PM-localized Sla1-positive patches and (E) Sla1-tdTom in PM-localized Mup1-positive patches intensity measurements of cells from different age categories (having 0-2, 3-5 or 5+ bud scars). Mean intensity measurements of age groups 3-5 and 5+ were normalized to those in the 0-2 age group. Data is reported as the statistical summary of the mean values ± standard deviation from at least three biological replicates. Means of individual biological replicates are represented with a circle, triangle, square, and diamond. At least 40 cells were analyzed per age group in each biological replicate. Statistical significance was assessed using paired t-tests. *, p<0.05; **, p<0.01; ns, not-significant. F) Representative micrographs of WGA-stained cells expressing Mup1-GFP and Sla1-tdTomato across different age groups. Shown are middle *z-*plane images, except for the WGA panels, which show maximum intensity projections. Arrowheads indicate examples of patches where Mup1 cargo colocalized with Sla1; shown also in insets within the merged image. Scale bar: 5 µm.

To generate intensity-time plots shown in Fig. 1I-K, a freehand line (line width = 1) was used to trace the movement of each patch and mean gray values were obtained from each time point (x-axis pixel on the kymograph). For each patch, individual timepoint intensity values were normalized to the track’s peak intensity value and were aligned to an arbitrary reference trajectory using a custom Python script. Intensity plots with multiple peaks (i.e., overlapping patches) were discarded from the analysis.

For the mother versus daughter analysis of Sla1 lifetime shown in Fig. 2, we selected budding cells and manually counted the bud scars of the larger (mother) cell. Daughter cells are located in bud scar-enriched areas due to the axial budding pattern of haploid cells (Chant & Pringle, 1995; Freifelder, 1960). This pattern helped to distinguish true budding cells from two adjacent cells with different cell sizes.

For the Mup1-GFP internalization and quinacrine staining intensity analyses, acquired z-stacks were transformed to maximum intensity z-projections using ImageJ. Whole-cell mean Mup1-GFP or quinacrine staining intensity was determined by measuring the intensity within the area obtained by drawing a circular outline around the outer edge of each cell using the ellipse tool. Cytosolic mean Mup1-GFP intensity (fluorescence inside the cell excluding the plasma membrane) was determined by measuring the intensity within the area obtained by drawing a circular outline a few pixels inward from the visible plasma membrane signal. For Mup1-GFP cargo and Sla1-tdTomato patch intensity analysis, first Sla1-tdTomato patches overlapping with Mup1-GFP patches were found by visual inspection of middle z-plane images, a square selection was then drawn tightly around the Sla1-tdTomato patch and intensity was measured for both the overlapping Mup1-GFP and Sla1-tdTomato patch. At least 40 cells for each age category were analyzed for each of the biological replicates.

### Statistical analyses

All experimental data are reported as a statistical summary of mean values from three biological replicates with standard deviation overlaid on a swarm plot of all individual measurements. Fluorescence intensity measurements for Mup1-GFP experiments (Fig. 3; and Fig. S2) were all normalized to age group 0-1 to account for variation in experimental setup. All statistical analyses were conducted using GraphPad Prism (GraphPad Software, Inc.) and a *P* value of less than 0.05 was considered statistically significant.

### Figure Preparation

For all figures, plots were created in GraphPad Prism (GraphPad Software, Inc.). Representative micrographs were generated by cropping individual cells from maximum-intensity projections of z-stacks, unless otherwise indicated, in ImageJ. Background fluorescence was subtracted from all cropped images with rolling ball radius of 50 pixels and pseudo-coloring was applied using lookup tables. All figures were assembled in Inkscape.

### Online Supplemental Material

**Table S1** contains a list of all yeast strains, plasmids and oligonucleotides used in the study. **Fig. S1** shows arrival dynamics of latter-arriving CME modules and CME protein abundance in young and older cells and complements data presented in Fig. 1. **Fig. S2** shows analysis of abundance of Mup1 in cells of various ages and complements the data shown in Fig. 3. **Fig. S3** shows representative micrographs, vacuole staining of young and old wildtype and *pma1-ts* cells under permissive and restrictive conditions, and Sla1 kymographs which complement data presented in Fig.4. **Fig. S4** shows that CME inhibition leads to loss vacuolar acidity. **Fig. S5** shows representative micrographs and Sla1 kymographs to complement the results shown in Fig. 6.

**Fig. 4.**
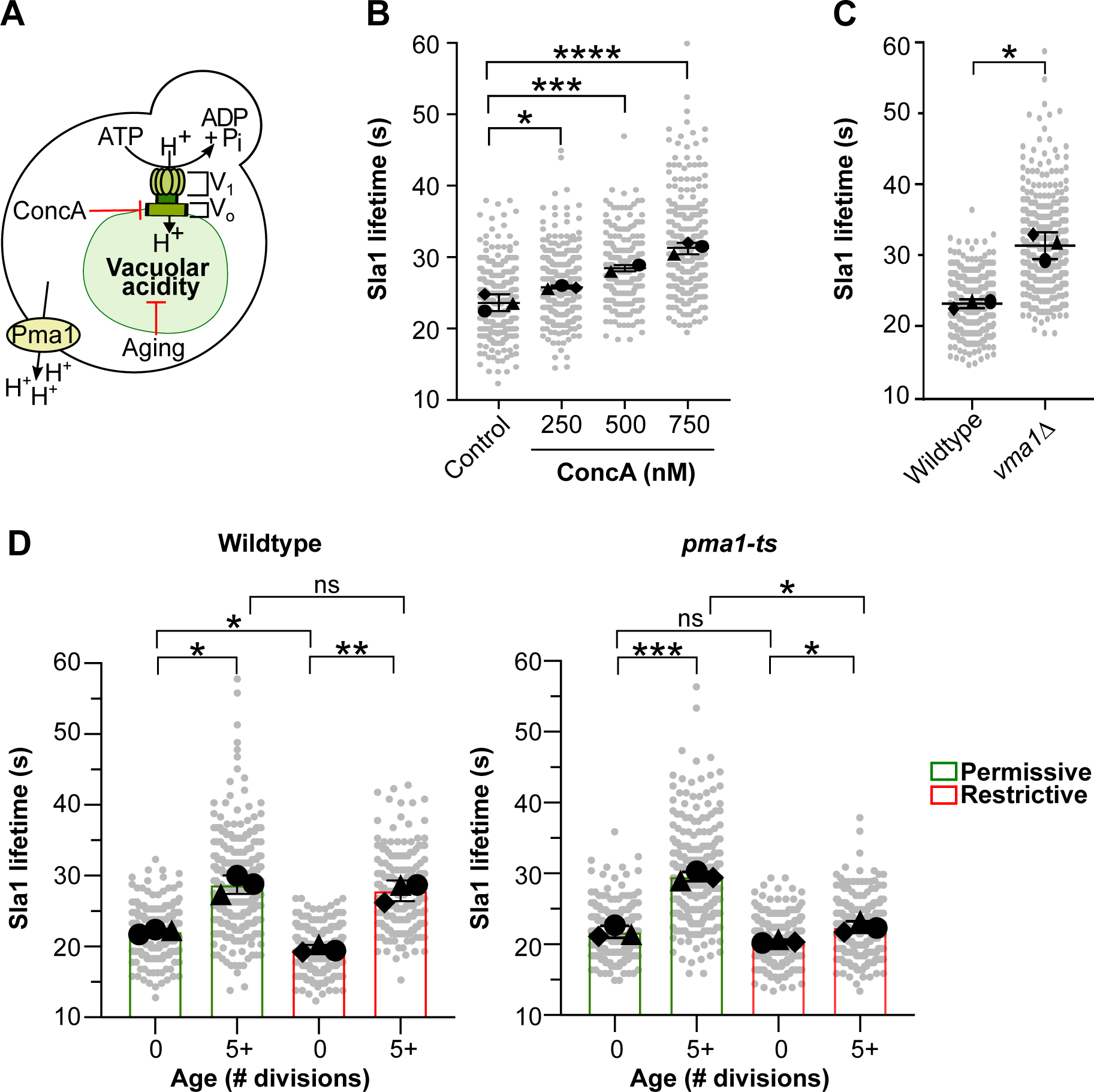
Alteration of vacuolar pH modulates normal progression of CME. A) Vacuolar pH in budding yeast can be altered through multiple mechanisms: V-ATPase activity can be inhibited using Concanamycin A (ConcA), via deletion of V-ATPase subunits, or through replicative aging during which the plasma membrane H^+^-ATPase, Pma1, accumulates in mother cells but not daughter cells. B) Quantification of Sla1 lifetimes in young (age 0-1) cells treated with increasing concentration of ConcA (250, 500 and 750 nM). C) Quantification of Sla1 lifetimes in young (age 0-1) wildtype and V-ATPase deficient, *vma1Δ* cells. D) Quantification of Sla1 lifetime in young (age 0-1) and old (age 5+) cells from wildtype and *pma1-ts* background under permissive (green bars) and restrictive (red bars) conditions. For all panels, superplots of Sla1 lifetime data are presented with a statistical summary of mean values ± standard deviation from three biological replicates (circle, triangle, diamond) overlayed to a swarm plot of individual measurements. At least 100 patches were measured from 10-20 cells for each condition, each age group, and each replicate. Statistical comparison was conducted using one-way ANOVA with Dunnett’s multiple comparison test for ConcA experiments (panel B), while paired t-test was used for the rest (panels C,D). *, p<0.05; **, p<0.01; ***, p<0.0001; ns, not-significant.

## RESULTS

### CME progression is delayed in replicatively-aging cells

We used real-time confocal microscopy and single-cell analysis to examine how the temporal dynamics of CME are altered during yeast replicative aging. CME progression depends on the proper function of several proteins categorized into distinct modules based on their shared roles (Fig. 1A). The lifetimes and recruitment dynamics of these CME proteins have been extensively characterized (Kaksonen et al., 2003, 2005). We focused on measuring patch lifetimes and recruitment dynamics of representative proteins from the early, coat, actin, myosin, and scission modules as a function of replicative age. The replicative age of individual unbudded mother cells was determined using the intensity of fluorophore-conjugated wheat germ agglutinin (WGA), a lectin that binds chitin, an N-acetylglucosamine polymer, enriched in bud scars (Guthrie & Fink, 2002). To distinguish between different age groups, we assumed a geometric distribution of replicative age within an asynchronous budding yeast population (Fig. 1B). Using this distribution, we defined virgin daughter cells (age 0 cells) as those belonging to the bottom 50% of WGA-staining intensities, age 1 cells as those belonging to the next 25% of the population, and so forth, up to the aged cells category (age 5+ cells) composed of cells belonging to the top 3% (cells with highest WGA staining intensities). We then generated kymographs along the plasma membrane of cells from each age group and quantified the lifetime of individual CME patches.

First, we first analyzed the lifetime of Sla1 in mother cells of different ages. Sla1 is a late-coat protein which has been used extensively as a marker of endocytic progression with a well characterized lifetime in wildtype and several endocytic mutants. We observed a clear increasing trend in the lifetimes of Sla1 patches with increasing replicative age (Fig. 1C). This trend reached statistical significance when comparing cells that have divided four times or more (age groups 4 and 5+) (age 4 bin: 29.2 ± 2.6 s; age 5+ bin: 32.6 ± 3.6 s) to virgin daughter cells (age 0) (23.1 ± 2.4 s).

To determine if this aging phenotype is present throughout the CME functional modules, we compared patch lifetimes and recruitment times relative to Sla1 of representative proteins from the five CME modules between virgin daughter and age 5+ cells using two-color live-cell fluorescence microscopy. First, to determine if the delayed progression of CME, shown by the prolonged Sla1 lifetime (Fig. 1C), was due to disruption of the recruitment dynamics of early CME proteins, we analyzed the lifetimes of Ede1, an early coat protein involved in site initiation in coordination with the yeast epsin homolog, Ent1 (Aguilar et al., 2003). We found that the lifetime of Ede1 was significantly longer in age 5+ cells demonstrating a difference of 13.6 ± 2.5 s relative to age 0 cells (Fig. 1D), corresponding to the prolonged Sla1 lifetime also observed in these cells (Fig. 1C). Our analysis also revealed that Sla1’s recruitment to Ede1 patches is delayed in age 5+ cells by 7.9 ± 1.5 s compared to age 0 cells (Fig. S1A). Next, we focused on representative proteins that are recruited after Sla1, namely Arp2 (actin nucleator), Sac6 (yeast fimbrin), Myo5 (yeast type I myosin), and Rvs167 (yeast amphiphysin). In contrast to Ede1, the lifetimes of the later-arriving CME modules were not different between age 0 and 5+ cells (Fig. 1E-H). This indicates that aging specifically affects the dynamics of the early steps of patch formation.

To support our patch lifetime results, we analyzed whether the recruitment dynamics, at the individual nascent-vesicle level, of later-arriving CME proteins differs between young and older cells. In wildtype cells, the coat module is recruited after early proteins form stable connections with the cargo. Then Sla1 reaches its peak intensity at the plasma membrane 15-20 seconds after recruitment (Kaksonen et al., 2005; Picco et al., 2015). Later-arriving CME proteins, including those in the actin, myosin and scission modules, are typically recruited once Sla1 reaches its peak intensity. We extracted intensity values as a function of time from individual Ede1, Sla1, Arp2 and Rvs167 patches from age 0 and 5+ cells. These intensity measurements were normalized to the respective protein’s peak intensity, and the resulting individual patch intensity plots were aligned at the time of peak intensity to generate average intensity profiles for each protein. Shown in Fig. 1I, the time required to reach peak intensity for Ede1 is approximately 40 seconds in age 5+ cells compared to 35 seconds observed in age 0 cells. Since Ede1’s recruitment phase takes longer in age 5+ cells, initiation of Sla1’s recruitment took approximately 30 seconds in age 5+ cells, compared to 20 seconds in age 0 cells. After peak recruitment, the total duration of the Sla1 intensity decline (indicative of vesicle internalization) is similar between the two age groups. These recruitment dynamics reflect the time of arrival of Sla1 (approx. 7.9 ± 1.5 s difference) to Ede1 patches shown above (Fig. S1A).

Focusing on the later-arriving CME modules, the recruitment profile of Arp2 did not differ significantly between the two age groups, with both requiring approximately 10 seconds to reach peak intensity thereby supporting the similar Arp2 lifetimes observed for the two age groups (Fig. 1J). The percentage of Sla1 intensity required to initiate Arp2 recruitment was also unchanged for the two age groups, with Arp2 recruitment starting as soon as Sla1 intensity reached approximately 50% of its peak intensity. However, overlaying the intensity plots of Sla1 and Arp2 from age 0-1 and 5+ cells revealed significant differences in Arp2 recruitment dynamics. Because Sla1 reaches peak intensity later in age 5+ cells, initiation of Arp2 recruitment was significantly delayed, taking approximately 20s compared to 10s in age 0-1 cells (Fig. S1B). Analysis of Rvs167 intensity profiles revealed recruitment dynamics similar to those of Arp2 (Fig. 1K, S1E). Overall, these results reinforce our previous observations that aging-related CME defects affect the dynamics of early modules while the later modules have relatively normal recruitment profiles and lifetimes.

To determine whether the altered lifetimes and recruitment profiles in aged cells are due to changes in the abundance of the studied CME proteins, we determined the replicative age of individual cells through manual annotation of their bud scars, and measured the whole-cell fluorescence intensity of CME markers for age 0-1 and 5+ cells. Our analysis revealed that abundance of Ede1, Sla1, Myo5 and Rvs167 proteins was unchanged in age 5+ cells compared to young cells (Fig. S1F,K). However, we observed a significant decrease in the abundance of the actin-related proteins, Sac6 and Arp2, in older cells (Fig. S1F,K). Our analyses also reveal a slight, though not significant, increase in the lifetime of actin-related proteins, Sac6 and Arp2, in older cells (Fig. 1E-F). This lifetime increase could be due to the lower abundance of these proteins in older cells resulting in reduced recruitment to the coat complex.

Collectively, our results reveal an aging-specific effect on the dynamics of CME progression. We show that the lifetimes and recruitment dynamics of the early and coat modules are delayed, which in turn affects the timing of the recruitment of the later-arriving modules. The observed changes in the early modules were not due to differences in protein abundance levels between young and older cells.

### Delayed CME progression is specific to mother cells and is not observed in daughter cells

Budding yeast undergoes asymmetric division, where aging-related factors are retained in the mother cell, leaving the daughter cell with rejuvenated replicative potential (Clay et al., 2014; Erjavec & Nyström, 2007; Higuchi-Sanabria et al., 2014; Shcheprova et al., 2008). During cell division, diffusion barriers form from septins and sphingolipids at the bud site (Fig. 2A). Given that mother cells retain most aging-associated cellular defects, we sought to determine whether the delayed CME progression observed in older cells was also mother-cell specific. We identified mother cells with medium to large buds (i.e. emerging daughters), and manually counted their WGA-stained bud scars, grouping them into three age categories: 0-2, 3-5 and 5+. We then quantified Sla1 patch lifetimes of individual mother cells and their buds (daughters), comparing the lifetimes based on the mother’s age group.

Consistent with our previous observations, we found a significant increase in Sla1 patch lifetimes with increasing replicative age in mother cells (Fig. 2B, representative micrographs shown in Fig. 2C). Specifically, age 0-2 mothers had an average Sla1 lifetime of 20.5 ± 0.9 s, age 3-5 mothers had an average Sla1 lifetime of 25.8 ± 1.9 s, and age 5+ mothers had an average Sla1 lifetime of 27.6 ± 1.5 s. In contrast, daughter cells displayed similar Sla1 patch lifetimes regardless of their mother’s replicative age. Additionally, we observed significant differences in Sla1 lifetimes between daughter cells and their mother across all age groups.

In conclusion, delayed CME progression is a replicative-age dependent and mother-cell specific phenotype that is not observed in newly formed daughter cells irrespective of their mother’s replicative age.

### Older cells internalize less endocytic cargo compared to young cells

Our results indicated that CME, specifically in the early cargo-dependent modules, is delayed in aging mother cells. To determine if this delay affects cargo internalization, we monitored the uptake of GFP-tagged Mup1, a high affinity methionine permease, as a CME cargo reporter in cells also expressing Sla1-tdTomato. Mup1 localizes to the plasma membrane in the absence of methionine, and is internalized under methionine excess or, similar to other PM transporters, in glucose starvation conditions (Fig. 3A) (Laidlaw et al., 2021). In both cases, Mup1 is ubiquitinated by the alpha-arrestin-ubiquitin ligase complex leading to its internalization via CME (Guiney et al., 2016; Ivashov et al., 2020). Upon methionine addition, Mup1 is rapidly internalized (Ivashov et al., 2020), while during glucose starvation, this process is much slower (Laidlaw et al., 2021), facilitating accurate quantification of the early steps of endocytosis.

We again categorized individual mother cells into three age groups (0-2, 3-5, and 5+) based on manual bud scar counts. Cultures were first grown in synthetic dextrose medium lacking methionine with 2% glucose (i.e. standard glucose medium), and z-stack images were acquired 10 minutes after switching to 2% raffinose medium (i.e. glucose starvation medium). For each age group, we measured the mean intensity of Mup1 of both whole cells and the cytosolic area (representing internalized cargo). We found decreased levels of cytosolic Mup1 in mother cells of ages 3-5 and 5+ compared to those from the age 0-2 group (Fig. 3B, representative micrographs shown in Fig. 3F). To determine the fraction of cargo internalized from the plasma membrane, we calculated the ratio of plasma membrane to cytosolic Mup1 (Fig. 3C). Older cells (from the 3-5 and 5+ age groups) had a higher membrane-to-cytosolic Mup1 ratio compared to age 0-2 cells, confirming reduced levels of internalized cargo in aging cells.

Next, we determined whether the amount of cargo associated with individual Sla1 patches differs in cells of varying replicative ages. We first identified Sla1 patches that colocalized with Mup1 cargo, and quantified the mean intensity of Mup1 at these patches. Similar to our other findings, older cells (from the 3-5 and 5+ age groups) showed a significant reduction in Mup1 cargo intensity at Sla1-positive patches compared to age 0-2 cells (Fig. 3D, representative micrographs shown in Fig. 3F). On the other hand, the amount of Sla1 recruited to each Mup1 cargo was not different between the age groups (Fig. 3E). These findings are consistent with previous reports showing that CME initiation is dependent on the amount of cargo recruited to the endocytic site, with low cargo levels leading to longer coat lifetimes (Carroll et al., 2012; Pedersen et al., 2020).

To determine if the decreased Mup1 internalization was due to differences in Mup1 protein levels between the studied age groups under normal glucose-replete conditions, we measured the overall Mup1 abundance in cells from each age group. Indeed, whole-cell Mup1 protein abundance was significantly lower in cells aged 3-5 and 5+ compared to young cells (from the age 0-2 group) (Fig. S2).

### Alteration of vacuolar pH delays CME progression

Vacuoles of aging mother cells have reduced acidity and this phenotype is evident as early as the 4th mitotic division (Hughes & Gottschling, 2012). Given our findings that early and coat module lifetimes are prolonged in mother cells that have divided 4 or more times (Figs. 1C-D), we next determined if this effect is a consequence or cause of altered vacuolar pH in aging cells. First, we treated wildtype cells with Concanamycin A (ConcA) (Fig. 4A), a specific V-ATPase inhibitor (Droese et al., 1993) at the indicated concentrations for 15 minutes, and analyzed Sla1 lifetimes in WGA-stained young cells only (0-1 age group). We used quinacrine, an amphipathic amine that accumulates in acidic compartments (Morano & Klionsky, 1994), to monitor vacuolar pH in all conditions. Our results show a concentration-dependent increase in Sla1 lifetimes in ConcA-treated cells compared to mock-treated control cells (Fig. 4B; Fig. S3A). Next, we examined Sla1 lifetimes in a *VMA1* deletion strain, *vma1Δ,* which lacks a subunit of the V_1_ subcomplex resulting in loss of vacuolar acidity regardless of a cell’s age (Forgac, 1999; J. Liu & Kane, 1996). In young *vma1Δ* cells, we observed significantly longer Sla1 lifetimes (Fig. 4C; Fig. S3B), similar to the effect observed in young cells treated with 750 nM ConcA. These results suggest that vacuolar pH plays a critical role in regulating CME dynamics.

To further investigate whether the observed effect on CME dynamics is V-ATPase-specific or depends on broader changes in cellular pH, we monitored Sla1 lifetimes in a temperature-sensitive *PMA1* loss-of-function mutant (*pma1-ts*). Pma1 has an asymmetric localization between aging mother cells and their daughters and has been shown to lead to increased vacuolar pH in older cells (Henderson et al., 2014). In the *pma1-ts* strain, under permissive conditions, aging mother cells have functional Pma1 resulting in a less acidic cytosolic pH, while under restrictive conditions, the loss of functional Pma1 reduces aging-associated loss of acidity (Fig. S3C). We quantified Sla1 lifetimes of young (age 0-1) and older (age 5+) *pma1-ts* and wildtype mother cells under both permissive and restrictive conditions (Fig. 4D, S3C). wildtype cells showed the expected Sla1 lifetime difference between young and older cells under both conditions, confirming that temperature changes alone don’t affect vacuolar pH. In *pma1-ts* cells, older mother cells exhibited longer Sla1 lifetimes under permissive conditions, similar to wildtype cells. However, under restrictive conditions, the difference between young and older *pma1-ts* cells was greatly reduced (Fig. 4D, S3C). Young *pma1-ts* cells showed no significant difference in Sla1 lifetimes between permissive and restrictive conditions. These results indicate that the temperature-sensitive *pma1* mutation predominantly affects aging mother cells, likely due to the preferential accumulation of Pma1 in these cells. Under restrictive conditions, older *pma1-ts* mother cells are unable to export protons, preserving vacuolar acidity during replicative aging. Taken together, our results show that CME dynamics are sensitive to changes in vacuolar pH during replicative aging through a mechanism linked to V-ATPase activity.

To explore if reduction of CME and subsequent retention of Pma1 in the plasma membrane of young cells leads to changes in vacuole acidity, we quantified vacuole pH of young cells with inhibited CME using genetic or chemical approaches. We used several CME mutants (*sla1Δ, cap1Δ* and *rvs167Δ*) that have been previously reported to have slower CME vesicle formation times relative to wildtype (Kaksonen et al., 2005), and quantified vacuole acidity using quinacrine staining. Shown in Fig. S4A, young *sla1Δ* cells demonstrate slight, yet significant, decrease in quinacrine staining intensity compared to wildtype cells, indicative of a decrease in vacuole acidity. In contrast, quinacrine staining intensity of young *cap1Δ* and *rvs167Δ* cells was not different from wildtype, potentially due to functional redundancy with *CAP2* and *RVS161* (Amatruda et al., 1990; Lombardi & Riezman, 2001).

Next, we used Latrunculin A (LatA), a toxin produced by *Latrunculia magnifica* that sequesters actin monomers and prevents their polymerization (Coué et al., 1987), that affects both actin cables and CME actin patches, and results in loss of polarity during yeast budding and blockage of CME (Kaksonen et al., 2003; Morton et al., 2000). We quantified quinacrine staining intensity in young cells under control and LatA-treatment conditions. LatA-treated young cells show a slight, yet significant, decrease in quinacrine intensity when compared to control cells (Fig. S4B), indicating a decrease in vacuole acidity. Collectively, these data show that reduction in CME dynamics during replicative aging could indeed promote retention of Pma1 at the plasma membrane of older cells, exacerbating the vacuole-mediated aging phenotypes.

### Restoring vacuolar acidity rejuvenates CME dynamics in aging cells

The reversible disassembly of V-ATPase in budding yeast is regulated by two factors, Oxr1 and the RAVE complex. The RAVE complex is a pro-assembly factor while Oxr1 is a disassembly factor that binds the V_1_ subcomplex of the V-ATPase following its dissociation from the V_o_ subcomplex (Fig. 5A) (Khan & Wilkens, 2024; Smardon et al., 2002). To further explore the role of vacuolar pH maintenance in CME dynamics, we investigated the effect of *OXR1* deletion. Oxr1 overexpression has been shown to reduce V-ATPase activity by promoting disassembly, while *OXR1* deletion (*oxr1Δ*) results in higher levels of assembled V-ATPase promoting vacuolar acidification (Klössel et al., 2024). Recent work has shown that Vma5, a subunit of the V_1_ subcomplex and specific binding partner of Oxr1, becomes increasingly cytosolic with increased replicative age suggesting that V-ATPase disassembly plays a role in vacuole pH dysregulation during aging (Hashmi & Kane, 2024). The *oxr1Δ* mutant is a longevity mutant with a 47.8% increase in replicative lifespan relative to wildtype (Hashmi & Kane, 2024), further highlighting the connection between vacuolar pH homeostasis and aging in budding yeast. We then asked whether Sla1 lifetimes between young and old cells differ in the *oxr1Δ* mutant compared to wildtype cells. Since the *oxr1Δ* mutant is expected to maintain higher V-ATPase activity (due to reduced disassembly) and vacuolar acidity during aging, we hypothesized that Sla1 lifetimes would remain largely unchanged between young and old cells in this mutant. As shown in Fig. 5A, we observed no significant difference in Sla1 lifetimes between young (age 0-1) and older cells (age 5+) in the *oxr1Δ* strain, in contrast to the lifetime difference observed between the two age groups in the wildtype strain.

**Fig. 5.**
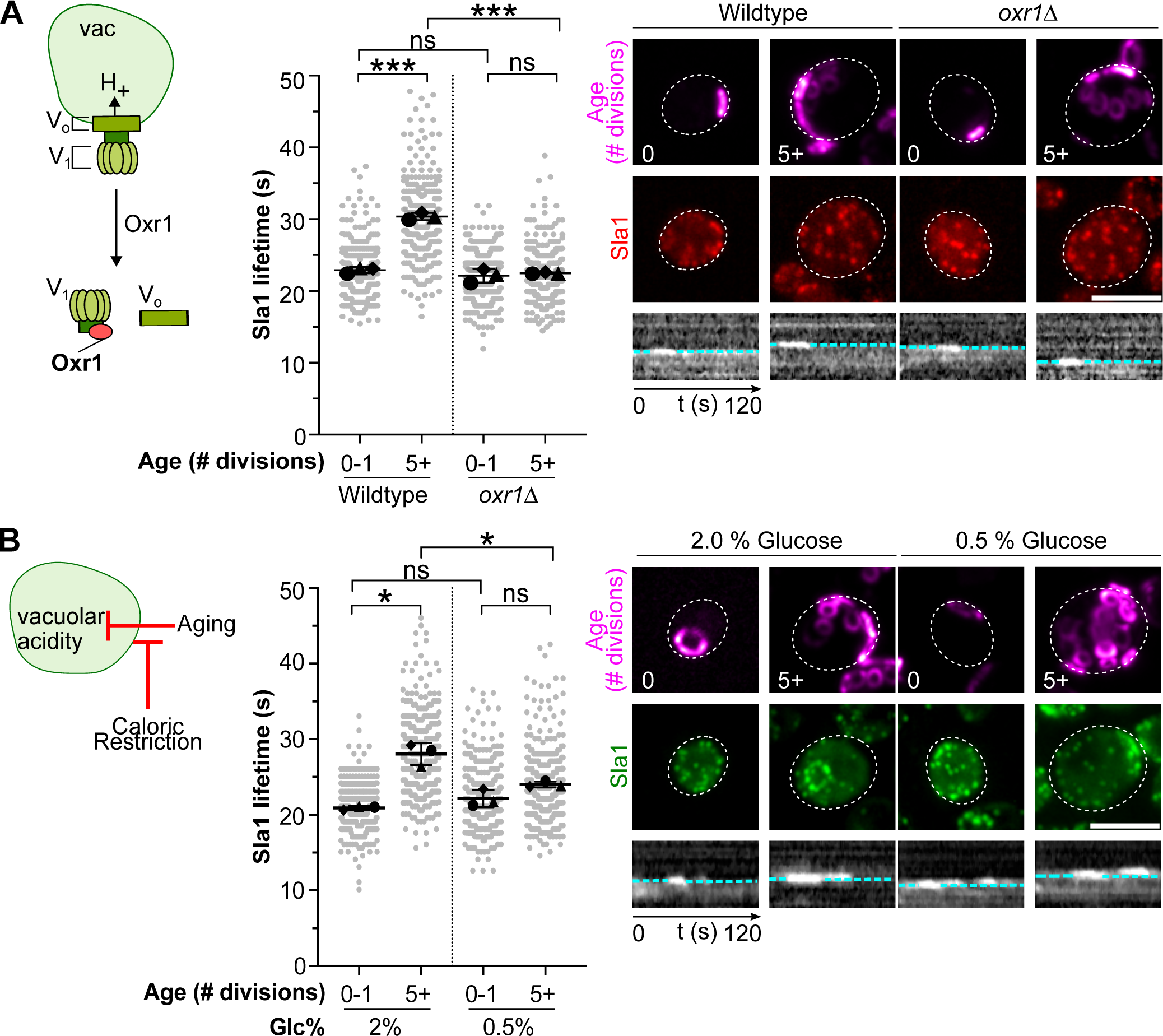
Maintenance of vacuolar acidity during replicative aging restores normal CME dynamics in older cells. A) Sla1 lifetime analysis in wildtype and *oxr1Δ* V-ATPase disassembly mutant. Left: Schematic diagram of the reversible yeast V-ATPase disassembly, regulated by the disassembly factor Oxr1, which binds specifically to the V_1_ subcomplex. Middle: Sla1 lifetime data were collected from young (age 0-1) and old (age 5+) wildtype and *oxr1Δ* cells. B) Quantification of Sla1 lifetimes in replicatively-aging cells under caloric restriction. Left: Schematic diagram of reversal of aging-linked vacuolar pH dysfunction by caloric restriction. Middle: Sla1 lifetime data were collected from young (age 0-1) and old (age 5+) wildtype cells grown in synthetic complete medium supplemented with either 2% or 0.5% glucose. For all experiments, Sla1 lifetime data are shown as mean values ± standard deviation from three biological replicates (circle, triangle, diamond), overlayed on a swarm plot of individual measurements. At least 100 Sla1 patches were quantified from 10-20 cells per strain for each age group and for each replicate. Statistical comparisons were done using a two-tailed paired t-test. *, p<0.05; **, p<0.01; ***, p<0.0001; ns, not-significant. Representative micrographs in A) and B): Maximum-intensity projections of z-stacks showing staining for age/bud scars (WGA) and Sla1-mCherry (in A) or Sla1-mNeonGreen (in B). Representative kymographs of Sla1 patches are shown under each micrograph. Plasma membrane is indicated by dashed lines, with the cytosol at the bottom. Scale bar: 5 µm.

Caloric restriction (CR) has been shown to extend lifespan in many eukaryotes by reversing numerous age-associated phenotypes, including the increase in vacuolar acidity by reducing V-ATPase disassembly in aging yeast cells (Hashmi & Kane, 2024; Lee et al., 1999; Lin et al., 2002). Since our results established a link between vacuolar pH and CME dynamics, we hypothesized that CR might restore a young cell-like CME phenotype by rejuvenating vacuolar acidity in aging cells. In budding yeast, CR is typically achieved by culturing cells in a nutritionally complete medium with reduced glucose (0.5% w/v compared to the standard 2%). As shown in Fig. 5B, under CR conditions, Sla1 lifetimes did not differ between young and old cells, while a significant difference was observed in standard medium. Furthermore, Sla1 lifetime in old cells grown under CR conditions was significantly shorter than in old cells grown in standard medium, indicating that CR mitigates the aging-related increase in Sla1 lifetimes. As expected, Sla1 lifetime in young cells was not affected by CR, since young cells already have an optimally-functioning acidic vacuole. These results suggest that CR rejuvenates CME dynamics in old cells by restoring vacuolar acidity.

Together, these results show that the maintenance of vacuolar acidity during replicative aging, achieved either through CR or genetic manipulation (*oxr1Δ*), restores CME dynamics to a more young cell-like state.

### Vacuolar pH regulates CME through TORC1-Npr1 signaling during replicative aging

The pH gradient between the cytosol and the vacuole is required for metabolite exchange between these two compartments. Vacuolar pH regulation is tightly linked to signaling complexes localized at the vacuolar membrane, with TORC1 acting as the key regulator of both anabolic and catabolic processes (Wullschleger et al., 2006). Previous studies have established that TORC1, through Rab-GTPases, responds to cellular nutrient levels in both yeast and mammalian cells (Bar-Peled et al., 2012; Panchaud et al., 2013). In yeast, downstream targets of TORC1 are involved in adaptation to cellular stress and cell growth (Barbet et al., 1996; Cherkasova & Hinnebusch, 2003). Other studies suggest that vacuolar pH, via a V-ATPase-dependent mechanism, negatively affects TORC1 activity. For example, deletion of the V-ATPase subunit Vma2 or induction of Pma1 expression using a doxycycline-repressible allele (*tetO7-PMA1*) both result in decreased phosphorylation of Sch9, the yeast homolog of the S6 kinase and key TORC1 target (Dechant et al., 2014). Given these findings, we explored whether the pH-dependent effect on CME is mediated by TORC1 signalling.

First, to determine whether TORC1 regulates CME dynamics during replicative aging, we quantified Sla1 lifetimes in young cells treated with the TORC1 inhibitor, rapamycin. We hypothesized that if vacuolar pH regulates CME dynamics through TORC1, inhibiting TORC1 in young cells would result in aged-cell-like Sla1 lifetimes regardless of their vacuolar pH status. Indeed, we observed a significant increase in Sla1 lifetimes in rapamycin-treated young cells compared to mock-treated control young cells (Fig. 6A, S5A).

**Fig. 6.**
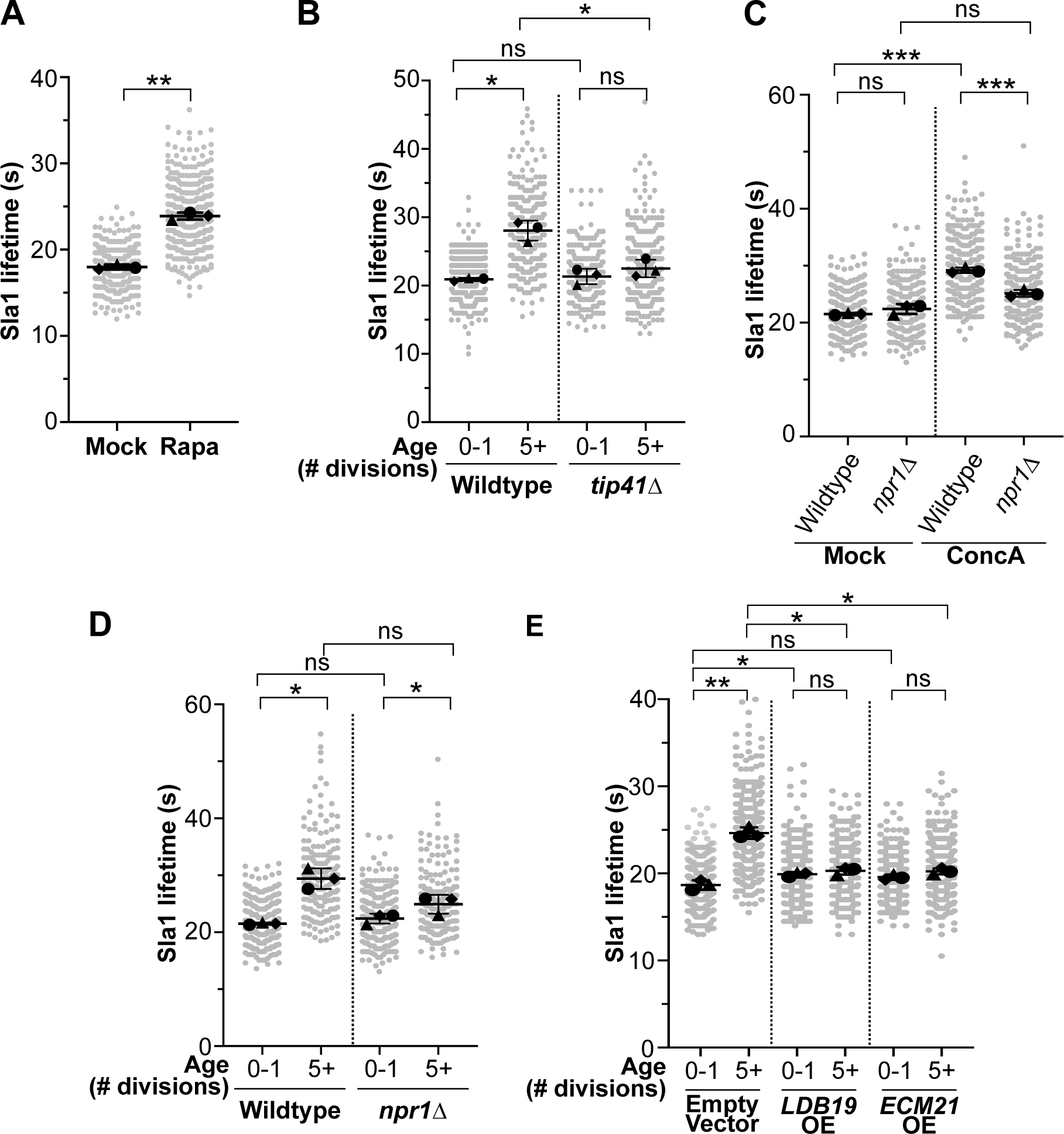
Vacuolar pH controls CME dynamics through TORC1-dependent signaling. A) Quantification of Sla1 lifetimes in young (age 0-1) wildtype cells treated with 200 ng/mL rapamycin for 2 hours. B) Quantification of Sla1 lifetimes in young (age 0-1) and old (age 5+) cells from wildtype and *tip41Δ* strain. C) Quantification of Sla1 lifetimes in young (age 0-1) wildtype and *npr1Δ* cells treated with 500 nM ConcA for 15 minutes. D) Quantification of Sla1 lifetimes in young (age 0-1) and old (age 5+) cells from wildtype and *npr1Δ* strain. E) Quantification of Sla1 lifetimes in young (age 0-1) and old (age 5+) wildtype cells over-expressing *LDB19* and *ECM21* arrestins. For all experiments, Sla1 lifetime data are shown as mean values ± standard deviation from three biological replicates (circle, triangle, diamond), overlayed on a swarm plot of individual measurements. At least 100 Sla1 patches were quantified from 10-20 cells per strain for each age group and for each replicate. Statistical comparisons were done using a two-tailed paired t-test. *, p<0.05; **, p<0.01; ***, p<0.0001; ns, not-significant.

Next, we assessed Sla1 lifetimes in a mutant harboring a deletion of *TIP41*, a negative regulator of TORC1 (Jacinto et al., 2001). We hypothesized that deletion of *TIP41* would counteract the inhibitory effect of low TORC1 activity on CME dynamics in high vacuolar pH conditions present in older cells. That is, in the *tip41Δ* strain, TORC1 remains active even under high vacuolar pH conditions. Consistent with our hypothesis, we observed no significant difference in Sla1 lifetimes between young and older *tip41Δ* cells, and both groups resembled young wildtype cells (Fig. 6B, S5A).

Previous studies have shown that yeast amino acid transporters, such as Gap1, Mup1 and Tat2, are regulated by the serine/threonine kinase Npr1, which is in turn under the control of TORC1 through phospho-inhibition (Craene et al., 2001; MacGurn et al., 2011; Schmidt et al., 1998). Npr1 inhibits alpha-arrestins, including Aly2/Art3 and Ldb19/Art1, which are responsible for the ubiquitin-mediated internalization of amino acid transporters (Ghaddar et al., 2014; Ivashov et al., 2020; O’Donnell et al., 2010). We observed that older cells internalize significantly less Mup1 under glucose-starvation conditions compared to young cells (Fig. 3), We hypothesized that in older cells with less acidic vacuolar pH, reduced TORC1 activity leads to loss of Npr1 inhibition, which in turn results in reduced activity of alpha-arrestins (ubiquitin ligase adaptors), and hence reduced levels of ubiquitinated cargo, impairing CME progression. To test this, we assessed CME dynamics in an *NPR1* deletion strain under ConcA treatment. Young *npr1Δ* cells showed no significant difference in Sla1 lifetimes between mock- and ConcA-treated conditions (Fig. 6C, S5C), in contrast to the longer Sla1 lifetimes observed in ConcA-treated wildtype cells. We also found that the difference in Sla1 lifetimes between young and older *npr1Δ* cells, while still significant, is less pronounced compared to wildtype cells (Fig. 6D, S5D).

We then questioned whether increasing the levels of Npr1-controlled alpha-arrestins can mitigate the CME progression defect observed in older cells exhibiting loss of TORC1-mediated Npr1 inhibition. The assumption here is that levels of non-phosphoinhibited alpha-arrestins would also increase upon overexpression of alpha-arrestins, facilitating normal cargo recruitment during CME. Sla1 lifetimes of older cells over-expressing Ldb19/Art1 from a high-copy (2µ) plasmid were significantly shorter than those of older cells carrying an empty vector (Fig. 6E, S5E). These results strongly suggest that the effect of high vacuolar pH on CME dynamics is mediated through TORC1 and alpha-arrestin signaling. Interestingly, over-expression of an Npr1-independent alpha-arrestin, Ecm21/Art2 (Savocco et al., 2019), has the same significant effect on Sla1 lifetimes (Fig. 6E, S5E), suggesting the existence of additional regulators besides Npr1.

Collectively, these results show that vacuolar pH modulates CME dynamics through TORC1-Npr1 signaling (Fig. 7). In young cells with normal acidic vacuolar pH, TORC1 inhibits Npr1, which in turn limits Art-Rsp5 complex activity required for cargo ubiquitination and CME progression. During replicative aging, dysfunctional vacuolar pH results in reduced TORC1 activity, leading to reduced phospho-inhibition of Npr1 and subsequent impairment of CME.

**Figure 7.**
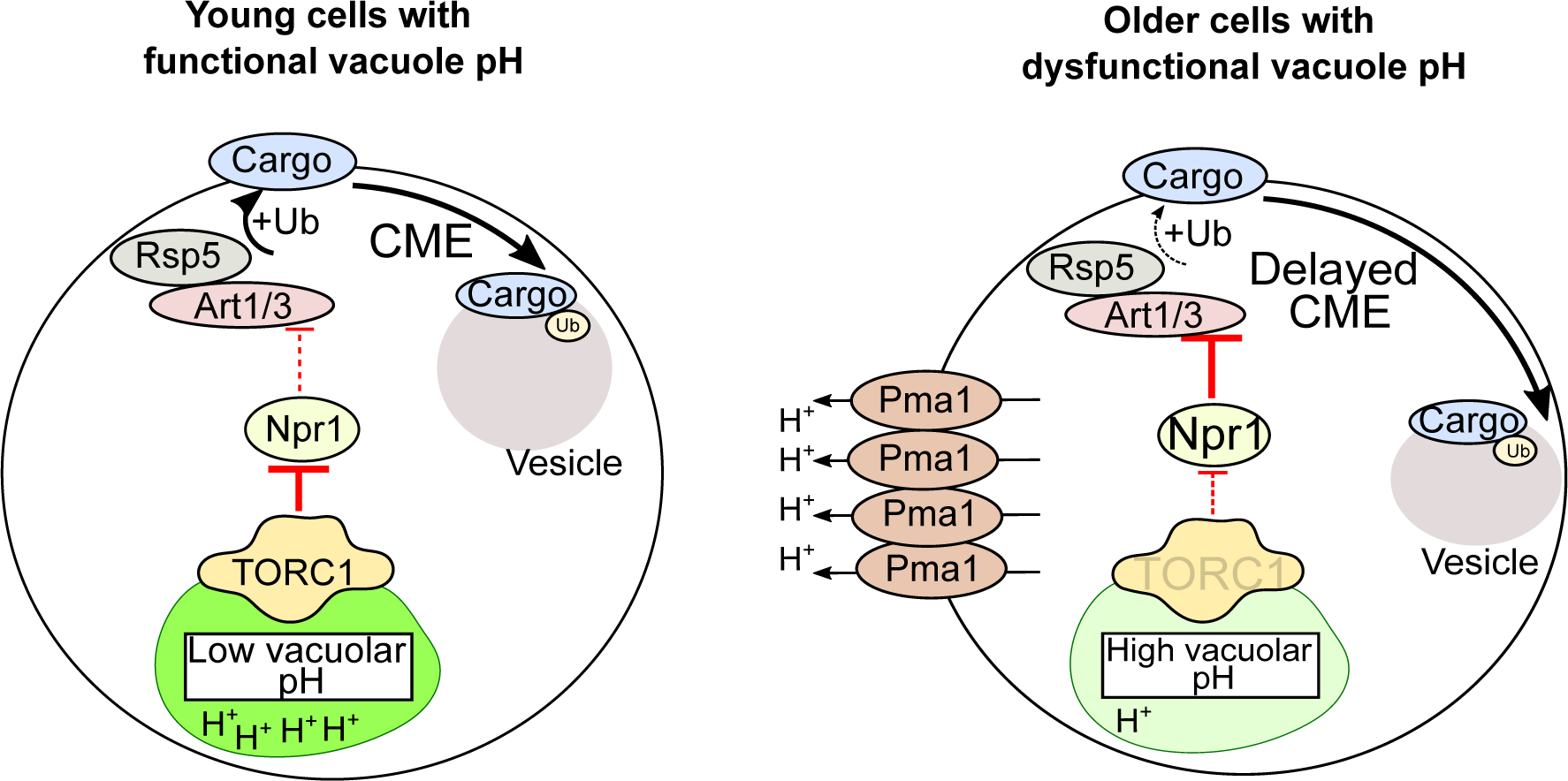
Proposed model of regulation of CME dynamics by TORC1 signaling in young and old cells. See main text for details.

## DISCUSSION

In this study, we aimed to understand how cellular aging impacts CME, a bioprocess critical for maintaining cellular homeostasis. We demonstrated that CME dynamics, particularly of the early and coat modules, are altered in cells at an early age. We also show that cargo internalization is reduced in older cells, and that these changes are age-specific and occur asymmetrically between older mothers and their daughter cells. These CME defects observed in older cells depend on V-ATPase function and are regulated by the TORC1 pathway.

### Aging-specific disruption of early steps of the vesicle formation process

Our initial focus was on understanding how CME is affected during replicative aging by analyzing the dynamics of representative proteins from the five CME modules in young and old cells. Our results reveal a significant trend where lifetimes of proteins in the early and coat modules are longer in older cells (indicating slower assembly of these modules), while later-arriving proteins have unchanged lifetimes, but show delayed arrival to the assembled coat module (Fig. 1D-G, Fig. S1B-E). CME has two functional phases, the variable and regular phase, which were defined based on the variability of CME-protein dynamics. Although Ede1 is a variable phase protein (Carroll et al., 2012; Stimpson et al., 2009), we found a clear difference between its lifetimes in young and old cells (Fig. 1D). As previously reported, prolonged Ede1 lifetimes are associated with stalled CME sites waiting for cargo accumulation before proceeding to the next, coat modules recruitment, phase (Pedersen et al., 2020). In our study, all Ede1 patches matured in the presence of Sla1, despite their variable lifetimes. This suggests that aging specifically affects CME progression from cargo binding to the early steps of the vesicle formation process.

Our results confirm a strong link between cargo availability and normal CME progression. In older cells, we observed a reduction in the amount of internalized and Sla1-colocalized Mup1, a high-affinity methionine and cysteine permease (Fig. 3B,D), along with prolonged Ede1 and Sla1 lifetimes (Fig. 1A,B). We also showed that the amount of Sla1 recruited to each Mup1-containing nascent vesicle is not significantly different between young and older cells (Fig. 3E). These findings indicate that cargo levels affect CME lifetime (i.e. progression), rather than the amount of CME protein recruited to the sites. Mup1 was chosen as the cargo of interest due to its experimental utility in observing different endocytic localizations (plasma membrane-bound, cytosolic and vacuolar). We also observed an overall reduction in Mup1 levels with age (Fig. S2). During replicative aging, cysteine accumulates in the cytosol due to failed transport to vacuoles with elevated pH, contributing to mitochondrial dysfunction (Chen et al., 2020; Hughes et al., 2020). Cysteine, a thiol-containing amino acid and sensor of the yeast sulfur amino acid pathway (Hansen & Johannesen, 2000), is also a substrate of Mup1. This raises the intriguing possibility that accumulating cysteine levels may be involved in regulating Mup1 expression during aging (Kosugi et al., 2001). Although direct evidence linking cysteine accumulation to Mup1 expression is limited, transcription factors, such as Met32 and Met4, known to regulate sulfur metabolism, also regulate Mup1 expression (Blaiseau et al., 1997; McIsaac et al., 2012). Notably, we only analyzed Mup1 internalization during aging; further research is thus needed to elucidate whether the observed reduction in Mup1 reflects a broader age-related decline in membrane transporter levels (see also below for discussion on TORC1 effects on translation), or could potentially be methionine- and/or cysteine-specific.

### Vacuolar pH and TORC1 signalling as key regulators of CME during aging

Our study highlights the critical role of vacuolar pH in regulating CME during replicative aging. We found that loss of vacuolar acidity in young cells results in CME dynamics similar to those seen in older cells (Fig. 4B,C). On the other hand, maintenance of vacuolar pH through CR or deletion of the V-ATPase disassembly factor, Oxr1, preserves normal CME dynamics in young and older cells (Fig. 5). However, the mechanism by which caloric restriction sustains vacuolar acidity during replicative aging remains unclear. V-ATPase assembly is glucose-sensitive, and it has been shown previously that CR reverses V-ATPase disassembly associated with replicative aging, by modulating abundance of Rav2, a V-ATPase pro-assembly factor (Hashmi & Kane, 2024). We also showed that loss of Pma1 function affects CME dynamics during replicative aging (Fig. 4D). Pma1 is the major yeast plasma membrane proton transporter and its asymmetric retention at the plasma membrane of aging mother cells has been described as a precursor to age-linked vacuolar pH alkalinization (Henderson et al., 2014). Loss of Pma1 function results in restoration of vacuolar acidity in aging mother cells through maintenance of the cytosolic pool of H^+^ available for vacuolar acidification. The CME dynamics of older *pma1-ts* cells under restrictive conditions (Fig. 4D) partially resembles those of their young counterparts which supports a vacuolar pH-dependent control of CME during aging. Importantly, Pma1 turnover at the plasma membrane is regulated by CME (Liu et al., 2006; Pizzirusso & Chang, 2004); our observed delay in CME (Fig. 1) and effect of CME disruption on vacuole acidity (Fig. S4) could thus exacerbate the accumulation of Pma1 at the plasma membrane of aging mother cells, further disrupting cellular homeostasis and contributing to the aging process.

Our results also show that TORC1 and Npr1 play a critical role in the regulation of CME dynamics during yeast replicative aging (Figs 6). Deletion of *NPR1* (a negative regulator of Ldb19/Art1 and Aly2/Art3 alpha arrestins), the function of which is regulated by TORC1, rescues the CME lifetime defects observed in ConcA-treated (V-ATPase-inhibited) young cells (Fig. 6C). While CME lifetimes in old *npr1Δ* cells remained slightly elevated compared to young cells (Fig. 6D), the difference was not as pronounced as in wildtype cells, suggesting that Npr1 plays a significant, though likely not exclusive, role in the aging-associated delay in CME. Consistently, increasing the levels of both Ldb19/Art1, an Npr1-dependent alpha arrestin, and Ecm21/Art2, an Npr1-independent alpha arrestin, in older cells resulted in CME lifetimes similar to those observed in young cells (Fig. 6E). Ldb19 and Ecm21 regulate CME of specific amino acid transporters (Guiney et al., 2016; Ivashov et al., 2020; Lee et al., 2019; Savocco et al., 2019); additional alpha arrestins regulate CME of other amino acid and hexose transporters (Becuwe et al., 2012; Llopis-Torregrosa et al., 2016; O’Donnell et al., 2015). It will be thus crucial to explore how additional regulatory proteins and alpha arrestins are affected by replicative aging. Additionally, TORC1 is a major regulator of translation, and reduction of TORC1 activity during aging could explain the observed reduction in whole-cell Mup1 abundance (Fig S2). The overall delay in CME progression during aging is thus likely a combination of reduced cargo levels due to global changes in protein translation, as well as reduced ubiquitination capacity of the alpha arrestin-Rsp5 pathway.

Collectively, our results suggest a model where the TORC1-Npr1 pathway controls the dynamics of cargo recognition and subsequent CME in a vacuolar pH-dependent manner (Fig. 7). In young cells where vacuolar acidity is optimal, TORC1 activity inhibits Npr1, thereby allowing alpha arrestins to promote cargo ubiquitination and CME progression. However, during aging, reduced vacuolar acidity downregulates TORC1 activity, resulting in reduced Npr1 phospho-inhibition and reduced alpha arrestin-mediated cargo ubiquitination. This leads to delays in CME progression, further impairing cellular function. Our findings provide insights into how vacuolar function and TORC1 signalling, both highly conserved across eukaryotes, influence CME during aging, and highlight a potential mechanism through which nutrient uptake and cellular homeostasis may decline over time, potentially contributing to aging-related pathologies.

## ACKNOWLEDGEMENTS

The authors thank Drs. D. Drubin (University of California, Berkeley), B. Andrews (University of Toronto), C. Boone (University of Toronto), and C. Antonescu (Toronto Metropolitan University) for access to reagents and equipment, and Dr. H. Friesen (University of Toronto) for technical support. We are grateful to Dr. H. Friesen and Dr. C. Antonescu for critical reading of the manuscript. This work was supported by Toronto Metropolitan University, the Faculty of Science, and a grant from Natural Sciences and Engineering Research Council of Canada (NSERC 03741-2021 to MMU).

## AUTHOR CONTRIBUTIONS

KA designed the project, carried out and analyzed experiments, prepared figures, and wrote the manuscript. JL and NG performed additional data analyses. SS and AS constructed strains and performed additional experiments. MMU designed and supervised the project, secured funding, and wrote the manuscript.

## CONFLICTS OF INTEREST

The authors declare no competing financial interests.

**Fig. S1. Arrival time, but not lifetime, of later modules is affected in older cells.**

A-E) Quantification of Ede1 duration before Sla1 arrival (A), and Sla1 duration before arrival of Arp2, Sac6, Myo5 and Rvs167 (B-E) in age 0-1 and 5+ cells. Data is reported as the statistical summary of the mean values ± standard deviation from three biological replicates (circle, triangle, diamond). Individual patch measurements are displayed as a swarm plot. At least 100 patches were quantified from 10-20 cells per strain for each age group and for each replicate. Statistical significance was assessed using Welch’s t-tests. *, p<0.05.

F-K) Quantification of mean fluorescence intensity of CME markers in young and older cells. Data is shown as mean ± standard deviation of mean intensity values from three biological replicates. For each replicate, at least 100 cells were analyzed for each age category. Paired t-test was conducted for statistical comparison. *, p<0.05; ***, p<0.0001; ns, not-significant.

**Fig. S2. Mup1 protein abundance decreases with age.**

(A) Quantitative analysis of Mup1 abundance in cells from different replicative age groups. Mean fluorescence intensity is used to represent Mup1 abundance. Data is reported as the statistical summary of the mean values ± standard deviation from three biological replicates. Individual measurements are displayed as a swarm plot. At least 40 cells were analyzed per age group for each replicate. Statistical significance was assessed using paired t-tests. *, p<0.05.

(B) Maximum-intensity z-stack projections of representative cells expressing Mup1-GFP are shown with staining for age/bud scars (WGA). Scale bar: 5 µm.

**Fig. S3. Loss of vacuolar pH regulation leads to longer Sla1 lifetimes.**

Maximum-intensity z-stack projections of representative cells are shown with staining for age/bud scars (WGA), vacuolar pH (quinacrine, panel A,C), and Sla1-mCherry (panel B). Representative kymographs of Sla1 are shown under the micrographs. Plasma membrane is indicated by dashed lines, with the cytosol at the bottom. Scale bar: 5 µm.

**Fig. S4 Inhibition of CME results in reduction of vacuole acidity.**

A) Quantification of vacuole acidity using quinacrine staining intensity in young cells from wildtype and CME mutant backgrounds, *sla1Δ, cap1Δ* and *rvs167Δ*.

B) Quantification of vacuole acidity using quinacrine staining intensity in young cells treated with 25 µM LatA.

For all experiments, superplots of quinacrine intensity data are shown as mean values ± standard deviation from three biological replicates (circle, triangle, diamond), overlayed on a swarm plot of individual measurements. 30 cells per strain (A) or per condition (B) were quantified for each replicate. Statistical comparisons were done using a two-tailed paired t-test. *, p<0.05; ns, not-significant.

Maximum-intensity z-stack projections of representative cells are shown with staining for age/bud scars (WGA), vacuolar pH (quinacrine), and Sac6-mScarlet-I (panel B). Scale bar: 5 µm

**Fig. S5. TORC1-dependent effect on Sla1 lifetimes in older cells.**

Maximum-intensity z-stack projections of representative cells are shown with staining for age/bud scars (WGA), vacuole (quinacrine, panel C), and Sla1-mNeonGreen (panels A,B,E), or Sla1-mCherry (panel D). Representative kymographs of Sla1 are shown under the micrographs. Plasma membrane is indicated by dashed lines, with the cytosol at the bottom. Scale bar: 5 µm.

## Supplementary Table

**Table S1. List of strains, plasmids and oligonucleotides used in the study.**

## REFERENCES

Aguilar, R. C., Watson, H. A., & Wendland, B. (2003). The Yeast Epsin Ent1 Is Recruited to Membranes through Multiple Independent Interactions. Journal of Biological Chemistry, 278(12), 10737–10743. 10.1074/jbc.M211622200

Amatruda, J. F., Cannon, J. F., Tatchell, K., Hug, C., & Cooper, J. A. (1990). Disruption of the actin cytoskeleton in yeast capping protein mutants. Nature, 344(6264), 352–354. 10.1038/344352a0

Babayev, E., & Duncan, F. E. (2022). Age-associated changes in cumulus cells and follicular fluid: The local oocyte microenvironment as a determinant of gamete quality. Biology of Reproduction, 106(2), 351–365. 10.1093/biolre/ioab241

Balguerie, A., Sivadon, P., Bonneu, M., & Aigle, M. (1999). Rvs167p, the budding yeast homolog of amphiphysin, colocalizes with actin patches. Journal of Cell Science, 112(15), 2529– 2537. 10.1242/jcs.112.15.2529

Barbet, N. C., Schneider, U., Helliwell, S. B., Stansfield, I., Tuite, M. F., & Hall, M. N. (1996). TOR controls translation initiation and early G1 progression in yeast. Molecular Biology of the Cell, 7(1), 25–42. 10.1091/mbc.7.1.25

Barker, S. L., Lee, L., Pierce, B. D., Maldonado-Báez, L., Drubin, D. G., & Wendland, B. (2007). Interaction of the Endocytic Scaffold Protein Pan1 with the Type I Myosins Contributes to the Late Stages of Endocytosis. Molecular Biology of the Cell, 18(8), 2893–2903. 10.1091/mbc.e07-05-0436

Bar-Peled, L., Schweitzer, L. D., Zoncu, R., & Sabatini, D. M. (2012). Ragulator Is a GEF for the Rag GTPases that Signal Amino Acid Levels to mTORC1. Cell, 150(6), 1196–1208. 10.1016/j.cell.2012.07.032

Becuwe, M., Vieira, N., Lara, D., Gomes-Rezende, J., Soares-Cunha, C., Casal, M., Haguenauer-Tsapis, R., Vincent, O., Paiva, S., & Léon, S. (2012). A molecular switch on an arrestin-like protein relays glucose signaling to transporter endocytosis. The Journal of Cell Biology, 196(2), 247–259. 10.1083/jcb.201109113

Blaiseau, P. L., Isnard, A. D., Surdin-Kerjan, Y., & Thomas, D. (1997). Met31p and Met32p, two related zinc finger proteins, are involved in transcriptional regulation of yeast sulfur amino acid metabolism. Molecular and Cellular Biology, 17(7), 3640–3648. 10.1128/MCB.17.7.3640

Carroll, S. Y., Stimpson, H. E. M., Weinberg, J., Toret, C. P., Sun, Y., & Drubin, D. G. (2012). Analysis of yeast endocytic site formation and maturation through a regulatory transition point. Molecular Biology of the Cell, 23(4), 657–668. 10.1091/mbc.e11-02-0108

Chant, J., & Pringle, J. R. (1995). Patterns of bud-site selection in the yeast Saccharomyces cerevisiae. The Journal of Cell Biology, 129(3), 751–765. 10.1083/jcb.129.3.751

Chen, K. L., Ven, T. N., Crane, M. M., Brunner, M. L. C., Pun, A. K., Helget, K. L., Brower, K., Chen, D. E., Doan, H., Dillard-Telm, J. D., Huynh, E., Feng, Y.-C., Yan, Z., Golubeva, A., Hsu, R. A., Knight, R., Levin, J., Mobasher, V., Muir, M., … Wasko, B. M. (2020). Loss of vacuolar acidity results in iron-sulfur cluster defects and divergent homeostatic responses during aging in Saccharomyces cerevisiae. GeroScience, 42(2), 749–764. 10.1007/s11357-020-00159-3

Cherkasova, V. A., & Hinnebusch, A. G. (2003). Translational control by TOR and TAP42 through dephosphorylation of eIF2α kinase GCN2. Genes & Development, 17(7), 859– 872. 10.1101/gad.1069003

Clay, L., Caudron, F., Denoth-Lippuner, A., Boettcher, B., Frei, S. B., Snapp, E. L., & Barral, Y. (2014). A sphingolipid-dependent diffusion barrier confines ER stress to the yeast mother cell. eLife, 3. 10.7554/eLife.01883

Coué, M., Brenner, S. L., Spector, I., & Korn, E. D. (1987). Inhibition of actin polymerization by latrunculin A. FEBS Letters, 213(2), 316–318. 10.1016/0014-5793(87)81513-2

Craene, J.-O. D., Soetens, O., & André, B. (2001). The Npr1 Kinase Controls Biosynthetic and Endocytic Sorting of the Yeast Gap1 Permease. Journal of Biological Chemistry, 276(47), 43939–43948. 10.1074/jbc.M102944200

Dechant, R., Saad, S., Ibáñez, A. J., & Peter, M. (2014). Cytosolic pH Regulates Cell Growth through Distinct GTPases, Arf1 and Gtr1, to Promote Ras/PKA and TORC1 Activity. Molecular Cell, 55(3), 409–421. 10.1016/j.molcel.2014.06.002

Droese, S., Bindseil, K. U., Bowman, E. J., Siebers, A., Zeeck, A., & Altendorf, K. (1993). Inhibitory effect of modified bafilomycins and concanamycins on P- and V-type adenosinetriphosphatases. Biochemistry, 32(15), 3902–3906. 10.1021/bi00066a008

Erjavec, N., & Nyström, T. (2007). Sir2p-dependent protein segregation gives rise to a superior reactive oxygen species management in the progeny of Saccharomyces cerevisiae. Proceedings of the National Academy of Sciences, 104(26), 10877–10881. 10.1073/pnas.0701634104

Feliciano, D., & Pietro, S. M. D. (2012). SLAC, a complex between Sla1 and Las17, regulates actin polymerization during clathrin-mediated endocytosis. Molecular Biology of the Cell, 23(21), 4256–4272. 10.1091/mbc.e11-12-1022

Fiore, P. P. D., & Zastrow, M. von. (2014). Endocytosis, Signaling, and Beyond. Cold Spring Harbor Perspectives in Biology, 6(8), a016865–a016865. 10.1101/cshperspect.a016865

Forgac, M. (1999). Structure and properties of the vacuolar (H+)-ATPases. The Journal of Biological Chemistry, 274(19), 12951–12954. 10.1074/jbc.274.19.12951

Freifelder, D. (1960). Bud position in Saccharomyces cerevisiae. Journal of Bacteriology, 80(4), 567–568. 10.1128/jb.80.4.567-568.1960

Garcia, C. K., Wilund, K., Arca, M., Zuliani, G., Fellin, R., Maioli, M., Calandra, S., Bertolini, S., Cossu, F., Grishin, N., Barnes, R., Cohen, J. C., & Hobbs, H. H. (2001). Autosomal Recessive Hypercholesterolemia Caused by Mutations in a Putative LDL Receptor Adaptor Protein. Science, 292(5520), 1394–1398. 10.1126/science.1060458

Ghaddar, K., Merhi, A., Saliba, E., Krammer, E.-M., Prévost, M., & André, B. (2014). Substrate-Induced Ubiquitylation and Endocytosis of Yeast Amino Acid Permeases. Molecular and Cellular Biology, 34(24), 4447–4463. 10.1128/MCB.00699-14

Gietz, R. D., & Schiestl, R. H. (2007). High-efficiency yeast transformation using the LiAc/SS carrier DNA/PEG method. Nature Protocols, 2(1), 31–34. 10.1038/nprot.2007.13

Graham, L. A., & Stevens, T. H. (1999). Assembly of the yeast vacuolar proton-translocating ATPase. Journal of Bioenergetics and Biomembranes, 31(1), 39–47. 10.1023/a:1005455411918

Guiney, E. L., Klecker, T., & Emr, S. D. (2016). Identification of the endocytic sorting signal recognized by the Art1-Rsp5 ubiquitin ligase complex. Molecular Biology of the Cell, 27(25), 4043–4054. 10.1091/mbc.E16-08-0570

Guthrie, C., & Fink, G. R. (2002). Part C. Guide to yeast genetics and molecular biology. In Methods in Enzymology. Academic Press.

Hansen, J., & Johannesen, P. F. (2000). Cysteine is essential for transcriptional regulation of the sulfur assimilation genes in Saccharomyces cerevisiae. Molecular and General Genetics MGG, 263(3), 535–542. 10.1007/s004380051199

Hashmi, F., & Kane, P. M. (2024). V-ATPase Disassembly at the Yeast Lysosome-Like Vacuole Is a Phenotypic Driver of Lysosome Dysfunction in Replicative Aging. bioRxiv, 2024.07.23.604825. 10.1101/2024.07.23.604825

Henderson, K. A., Hughes, A. L., & Gottschling, D. E. (2014). Mother-daughter asymmetry of pH underlies aging and rejuvenation in yeast. eLife, 3. 10.7554/eLife.03504

Higuchi-Sanabria, R., Pernice, W. M. A., Vevea, J. D., Wolken, D. M. A., Boldogh, I. R., & Pon, L. A. (2014). Role of asymmetric cell division in lifespan control in Saccharomyces cerevisiae. FEMS Yeast Research, 14(8), 1133–1146. 10.1111/1567-1364.12216

Hou, Y., Dan, X., Babbar, M., Wei, Y., Hasselbalch, S. G., Croteau, D. L., & Bohr, V. A. (2019). Ageing as a risk factor for neurodegenerative disease. Nature Reviews Neurology, 15(10), 565–581. 10.1038/s41582-019-0244-7

Hughes, A., & Gottschling, D. (2012). An early age increase in vacuolar pH limits mitochondrial function and lifespan in yeast. Nature, 492(7428), 261–265. 10.1038/nature11654

Hughes, C. E., Coody, T. K., Jeong, M.-Y., Berg, J. A., Winge, D. R., & Hughes, A. L. (2020). Cysteine Toxicity Drives Age-Related Mitochondrial Decline by Altering Iron Homeostasis. Cell, 180(2), 296–310.e18. 10.1016/j.cell.2019.12.035

Idrissi, F.-Z., Wolf, B. L., & Geli, M. I. (2002). Cofilin, But Not Profilin, Is Required for Myosin-I-Induced Actin Polymerization and the Endocytic Uptake in Yeast. Molecular Biology of the Cell, 13(11), 4074–4087. 10.1091/mbc.02-04-0052

Ivashov, V., Zimmer, J., Schwabl, S., Kahlhofer, J., Weys, S., Gstir, R., Jakschitz, T., Kremser, L., Bonn, G. K., Lindner, H., Huber, L. A., Leon, S., Schmidt, O., & Teis, D. (2020). Complementary α-arrestin-ubiquitin ligase complexes control nutrient transporter endocytosis in response to amino acids. eLife, 9. 10.7554/eLife.58246

Jacinto, E., Guo, B., Arndt, K. T., Schmelzle, T., & Hall, M. N. (2001). TIP41 Interacts with TAP42 and Negatively Regulates the TOR Signaling Pathway. Molecular Cell, 8(5), 1017– 1026. 10.1016/S1097-2765(01)00386-0

Janssens, G., & Veenhoff, L. (2016). Evidence for the hallmarks of human aging in replicatively aging yeast. Microbial Cell, 3(7), 263–274. 10.15698/mic2016.07.510

Kaksonen, M., Sun, Y., & Drubin, D. G. (2003). A Pathway for Association of Receptors, Adaptors, and Actin during Endocytic Internalization. Cell, 115(4), 475–487. 10.1016/S0092-8674(03)00883-3

Kaksonen, M., Toret, C. P., & Drubin, D. G. (2005). A modular design for the clathrin- and actin-mediated endocytosis machinery. Cell, 123(2), 305–320. 10.1016/j.cell.2005.09.024

Khan, M. M., & Wilkens, S. (2024). Molecular mechanism of Oxr1p mediated disassembly of yeast V-ATPase. EMBO Reports, 25(5), 2323–2347. 10.1038/s44319-024-00126-5

Klössel, S., Zhu, Y., Amado, L., Bisinski, D. D., Ruta, J., Liu, F., & Montoro, A. G. (2024). Yeast TLDc domain proteins regulate assembly state and subcellular localization of the V-ATPase. The EMBO Journal, 43(9), 1870–1897. 10.1038/s44318-024-00097-2

Kosugi, A., Koizumi, Y., Yanagida, F., & Udaka, S. (2001). MUP1, high affinity methionine permease, is involved in cysteine uptake by Saccharomyces cerevisiae. Bioscience, Biotechnology, and Biochemistry, 65(3), 728–731. 10.1271/bbb.65.728

Laidlaw, K. M. E., Bisinski, D. D., Shashkova, S., Paine, K. M., Veillon, M. A., Leake, M. C., & MacDonald, C. (2021). A glucose-starvation response governs endocytic trafficking and eisosomal retention of surface cargoes in budding yeast. Journal of Cell Science, 134(2). 10.1242/jcs.257733

Lee, C. K., Klopp, R. G., Weindruch, R., & Prolla, T. A. (1999). Gene expression profile of aging and its retardation by caloric restriction. *Science (New York*, N.Y.), 285(5432), 1390–1393. 10.1126/science.285.5432.1390

Lee, S., Ho, H.-C., Tumolo, J. M., Hsu, P.-C., & MacGurn, J. A. (2019). Methionine triggers Ppz-mediated dephosphorylation of Art1 to promote cargo-specific endocytosis. The Journal of Cell Biology, 218(3), 977–992. 10.1083/jcb.201712144

Li, S. C., & Kane, P. M. (2009). The yeast lysosome-like vacuole: Endpoint and crossroads. Biochimica et Biophysica Acta (BBA) - Molecular Cell Research, 1793(4), 650–663. 10.1016/j.bbamcr.2008.08.003

Lin, C. H., MacGurn, J. A., Chu, T., Stefan, C. J., & Emr, S. D. (2008). Arrestin-Related Ubiquitin-Ligase Adaptors Regulate Endocytosis and Protein Turnover at the Cell Surface. Cell, 135(4), 714–725. 10.1016/j.cell.2008.09.025

Lin, S.-J., Kaeberlein, M., Andalis, A. A., Sturtz, L. A., Defossez, P.-A., Culotta, V. C., Fink, G. R., & Guarente, L. (2002). Calorie restriction extends Saccharomyces cerevisiae lifespan by increasing respiration. Nature, 418(6895), 344–348. 10.1038/nature00829

Liu, J., & Kane, P. M. (1996). Mutational Analysis of the Catalytic Subunit of the Yeast Vacuolar Proton-Translocating ATPase. Biochemistry, 35(33), 10938–10948. 10.1021/bi9608065

Liu, P., Young, T. Z., & Acar, M. (2015). Yeast Replicator: A High-Throughput Multiplexed Microfluidics Platform for Automated Measurements of Single-Cell Aging. Cell Reports, 13(3), 634–644. 10.1016/j.celrep.2015.09.012

Liu, Y., Sitaraman, S., & Chang, A. (2006). Multiple Degradation Pathways for Misfolded Mutants of the Yeast Plasma Membrane ATPase, PMA1*. Journal of Biological Chemistry, 281(42), 31457–31466. 10.1016/S0021-9258(19)84058-9

Llopis-Torregrosa, V., Ferri-Blázquez, A., Adam-Artigues, A., Deffontaines, E., van Heusden, G. P. H., & Yenush, L. (2016). Regulation of the Yeast Hxt6 Hexose Transporter by the Rod1 α-Arrestin, the Snf1 Protein Kinase, and the Bmh2 14-3-3 Protein*. Journal of Biological Chemistry, 291(29), 14973–14985. 10.1074/jbc.M116.733923

Lombardi, R., & Riezman, H. (2001). Rvs161p and Rvs167p, the Two Yeast Amphiphysin Homologs, Function Together in Vivo. Journal of Biological Chemistry, 276(8), 6016– 6022. 10.1074/jbc.M008735200

Longo, V. D., Shadel, G. S., Kaeberlein, M., & Kennedy, B. (2012). Replicative and Chronological Aging in Saccharomyces cerevisiae. Cell Metabolism, 16(1), 18–31. 10.1016/j.cmet.2012.06.002

Lu, R., Drubin, D. G., & Sun, Y. (2016). Clathrin-mediated endocytosis in budding yeast at a glance. Journal of Cell Science, 129(8), 1531–1536. 10.1242/jcs.182303

MacDiarmid, C. W., Gaither, L. A., & Eide, D. (2000). Zinc transporters that regulate vacuolar zinc storage in Saccharomyces cerevisiae. The EMBO Journal, 19(12), 2845–2855. 10.1093/emboj/19.12.2845

MacGurn, J. A., Hsu, P.-C., Smolka, M. B., & Emr, S. D. (2011). TORC1 Regulates Endocytosis via Npr1-Mediated Phosphoinhibition of a Ubiquitin Ligase Adaptor. Cell, 147(5), 1104– 1117. 10.1016/j.cell.2011.09.054

Magtanong, L., Ho, C. H., Barker, S. L., Jiao, W., Baryshnikova, A., Bahr, S., Smith, A. M., Heisler, L. E., Choy, J. S., Kuzmin, E., Andrusiak, K., Kobylianski, A., Li, Z., Costanzo, M., Basrai, M. A., Giaever, G., Nislow, C., Andrews, B., & Boone, C. (2011). Dosage suppression genetic interaction networks enhance functional wiring diagrams of the cell. Nature Biotechnology, 29(6), 505–511. 10.1038/nbt.1855

Maldonado-Báez, L., Dores, M. R., Perkins, E. M., Drivas, T. G., Hicke, L., & Wendland, B. (2008). Interaction between Epsin/Yap180 Adaptors and the Scaffolds Ede1/Pan1 Is Required for Endocytosis. Molecular Biology of the Cell, 19(7), 2936–2948. 10.1091/mbc.e07-10-1019

Mattiazzi Usaj, M., Sahin, N., Friesen, H., Pons, C., Usaj, M., Masinas, M. P. D., Shuteriqi, E., Shkurin, A., Aloy, P., Morris, Q., Boone, C., & Andrews, B. J. (2020). Systematic genetics and single-cell imaging reveal widespread morphological pleiotropy and cell-to-cell variability. Molecular Systems Biology, 16(2), 1–27. 10.15252/msb.20199243

McCormick, M. A., Delaney, J. R., Tsuchiya, M., Tsuchiyama, S., Shemorry, A., Sim, S., Chou, A. C.-Z., Ahmed, U., Carr, D., Murakami, C. J., Schleit, J., Sutphin, G. L., Wasko, B. M., Bennett, C. F., Wang, A. M., Olsen, B., Beyer, R. P., Bammler, T. K., Prunkard, D., … Kennedy, B. K. (2015). A Comprehensive Analysis of Replicative Lifespan in 4,698 Single-Gene Deletion Strains Uncovers Conserved Mechanisms of Aging. Cell Metabolism, 22(5), 895–906. 10.1016/j.cmet.2015.09.008

McIsaac, R. S., Petti, A. A., Bussemaker, H. J., & Botstein, D. (2012). Perturbation-based analysis and modeling of combinatorial regulation in the yeast sulfur assimilation pathway. Molecular Biology of the Cell, 23(15), 2993–3007. 10.1091/mbc.E12-03-0232

Mellman, I., & Yarden, Y. (2013). Endocytosis and Cancer. Cold Spring Harbor Perspectives in Biology, 5(12), a016949–a016949. 10.1101/cshperspect.a016949

Miseta, A., Kellermayer, R., Aiello, D. P., Fu, L., & Bedwell, D. M. (1999). The vacuolar Ca2+/H+ exchanger Vcx1p/Hum1p tightly controls cytosolic Ca2+ levels in S. cerevisiae. FEBS Letters, 451(2), 132–136. 10.1016/s0014-5793(99)00519-0

Morano, K. A., & Klionsky, D. J. (1994). Differential effects of compartment deacidification on the targeting of membrane and soluble proteins to the vacuole in yeast. Journal of Cell Science, 107(10), 2813–2824. 10.1242/jcs.107.10.2813

Mortimer, R. K., & Johnston, J. R. (1959). Life Span of Individual Yeast Cells. Nature, 183(4677), 1751–1752. 10.1038/1831751a0

Morton, W. M., Ayscough, K. R., & McLaughlin, P. J. (2000). Latrunculin alters the actin-monomer subunit interface to prevent polymerization. Nature Cell Biology, 2(6), 376–378. 10.1038/35014075

Myers, M. D., Ryazantsev, S., Hicke, L., & Payne, G. S. (2016). Calmodulin Promotes N-BAR Domain-Mediated Membrane Constriction and Endocytosis. Developmental Cell, 37(2), 162–173. 10.1016/j.devcel.2016.03.012

O’Donnell, A. F., Apffel, A., Gardner, R. G., & Cyert, M. S. (2010). α-Arrestins Aly1 and Aly2 Regulate Intracellular Trafficking in Response to Nutrient Signaling. Molecular Biology of the Cell, 21(20), 3552–3566. 10.1091/mbc.e10-07-0636

O’Donnell, A. F., McCartney, R. R., Chandrashekarappa, D. G., Zhang, B. B., Thorner, J., & Schmidt, M. C. (2015). 2-Deoxyglucose impairs Saccharomyces cerevisiae growth by stimulating Snf1-regulated and α-arrestin-mediated trafficking of hexose transporters 1 and 3. Molecular and Cellular Biology, 35(6), 939–955. 10.1128/MCB.01183-14

Okreglak, V., & Drubin, D. G. (2007). Cofilin recruitment and function during actin-mediated endocytosis dictated by actin nucleotide state. The Journal of Cell Biology, 178(7), 1251– 1264. 10.1083/jcb.200703092

Palmer, S. E., Rooij, I. I. S., Marklew, C. J., Allwood, E. G., Mishra, R., Johnson, S., Goldberg, M. W., & Ayscough, K. R. (2015). A Dynamin-Actin Interaction Is Required for Vesicle Scission during Endocytosis in Yeast. Current Biology, 25(7), 868–878. 10.1016/j.cub.2015.01.061

Panchaud, N., Péli-Gulli, M.-P., & Virgilio, C. D. (2013). Amino Acid Deprivation Inhibits TORC1 Through a GTPase-Activating Protein Complex for the Rag Family GTPase Gtr1. Science Signaling, 6(277), ra42. 10.1126/scisignal.2004112

Pedersen, R. T. A., & Drubin, D. G. (2019). Type I myosins anchor actin assembly to the plasma membrane during clathrin-mediated endocytosis. Journal of Cell Biology, 218(4), 1138– 1147. 10.1083/jcb.201810005

Pedersen, R. T. A., Hassinger, J. E., Marchando, P., & Drubin, D. G. (2020). Spatial regulation of clathrin-mediated endocytosis through position-dependent site maturation. Journal of Cell Biology, 219(11). 10.1083/jcb.202002160

Peng, Y., Grassart, A., Lu, R., Wong, C. C. L., Yates, J., Barnes, G., & Drubin, D. G. (2015). Casein Kinase 1 Promotes Initiation of Clathrin-Mediated Endocytosis. Developmental Cell, 32(2), 231–240. 10.1016/j.devcel.2014.11.014

Picco, A., Mund, M., Ries, J., Nédélec, F., & Kaksonen, M. (2015). Visualizing the functional architecture of the endocytic machinery. eLife, 4. 10.7554/eLife.04535

Pizzirusso, M., & Chang, A. (2004). Ubiquitin-mediated targeting of a mutant plasma membrane ATPase, Pma1-7, to the endosomal/vacuolar system in yeast. Molecular Biology of the Cell, 15(5), 2401–2409. 10.1091/mbc.e03-10-0727

Qu, Y., Meng, B., Cai, S., Yang, B., He, Y., Fu, C., Li, X., Li, P., Cao, Z., Mao, X., Teng, W., & Shi, S. (2024). Apoptotic metabolites ameliorate bone aging phenotypes via TCOF1/FLVCR1-mediated mitochondrial homeostasis. Journal of Nanobiotechnology, 22(1), 549. 10.1186/s12951-024-02820-x

Russnak, R., Konczal, D., & McIntire, S. L. (2001). A family of yeast proteins mediating bidirectional vacuolar amino acid transport. The Journal of Biological Chemistry, 276(26), 23849–23857. 10.1074/jbc.M008028200

Savocco, J., Nootens, S., Afokpa, W., Bausart, M., Chen, X., Villers, J., Renard, H.-F., Prévost, M., Wattiez, R., & Morsomme, P. (2019). Yeast α-arrestin Art2 is the key regulator of ubiquitylation-dependent endocytosis of plasma membrane vitamin B1 transporters. PLoS Biology, 17(10), e3000512.

Schmidt, A., Beck, T., Koller, A., Kunz, J., & Hall, M. N. (1998). The TOR nutrient signalling pathway phosphorylates NPR1 and inhibits turnover of the tryptophan permease. The EMBO Journal, 17(23), 6924–6931. 10.1093/emboj/17.23.6924

Schneider, C. A., Rasband, W. S., & Eliceiri, K. W. (2012). NIH Image to ImageJ: 25 years of image analysis. Nature Methods, 9(7), 671–675. 10.1038/nmeth.2089

Serrano, R. (1978). Characterization of the plasma membrane ATPase of Saccharomyces cerevisiae. Molecular and Cellular Biochemistry, 22(1), 51–63. 10.1007/BF00241470

Serrano, R., Kielland-Brandt, M. C., & Fink, G. R. (1986). Yeast plasma membrane ATPase is essential for growth and has homology with (Na+ + K+), K+- and Ca2+-ATPases. Nature, 319(6055), 689–693. 10.1038/319689a0

Shcheprova, Z., Baldi, S., Frei, S. B., Gonnet, G., & Barral, Y. (2008). A mechanism for asymmetric segregation of age during yeast budding. Nature, 454(7205), 728–734. 10.1038/nature07212

Smardon, A. M., Tarsio, M., & Kane, P. M. (2002). The RAVE Complex Is Essential for Stable Assembly of the Yeast V-ATPase *. Journal of Biological Chemistry, 277(16), 13831– 13839. 10.1074/jbc.M200682200

Stimpson, H. E. M., Toret, C. P., Cheng, A. T., Pauly, B. S., & Drubin, D. G. (2009). Early-Arriving Syp1p and Ede1p Function in Endocytic Site Placement and Formation in Budding Yeast. Molecular Biology of the Cell, 20(22), 4640–4651. 10.1091/mbc.e09-05-0429

Sun, Y., Leong, N. T., Wong, T., & Drubin, D. G. (2015). A Pan1/End3/Sla1 complex links Arp2/3-mediated actin assembly to sites of clathrin-mediated endocytosis. Molecular Biology of the Cell, 26(21), 3841–3856. 10.1091/mbc.E15-04-0252

Sun, Y., Martin, A. C., & Drubin, D. G. (2006). Endocytic Internalization in Budding Yeast Requires Coordinated Actin Nucleation and Myosin Motor Activity. Developmental Cell, 11(1), 33–46. 10.1016/j.devcel.2006.05.008

Tessneer, K. L., Pasula, S., Cai, X., Dong, Y., Liu, X., Yu, L., Hahn, S., McManus, J., Chen, Y., Chang, B., & Chen, H. (2013). Endocytic Adaptor Protein Epsin Is Elevated in Prostate Cancer and Required for Cancer Progression. ISRN Oncology, 2013, 1–8. 10.1155/2013/420597

Tolsma, T. O., Cuevas, L. M., & Pietro, S. M. D. (2018). The Sla1 adaptor-clathrin interaction regulates coat formation and progression of endocytosis. Traffic, 19(6), 446–462. 10.1111/tra.12563

Tolsma, T. O., Febvre, H. P., Olson, D. M., & Di Pietro, S. M. (2020). Cargo-mediated recruitment of the endocytic adaptor protein Sla1 in S. cerevisiae. Journal of Cell Science, 133(19). 10.1242/jcs.247684

Wullschleger, S., Loewith, R., & Hall, M. N. (2006). TOR Signaling in Growth and Metabolism. Cell, 124(3), 471–484. 10.1016/j.cell.2006.01.016

Youn, J.-Y., Friesen, H., Kishimoto, T., Henne, W. M., Kurat, C. F., Ye, W., Ceccarelli, D. F., Sicheri, F., Kohlwein, S. D., McMahon, H. T., & Andrews, B. J. (2010). Dissecting BAR Domain Function in the Yeast Amphiphysins Rvs161 and Rvs167 during Endocytosis. Molecular Biology of the Cell, 21(17), 3054–3069. 10.1091/mbc.e10-03-0181

