## Supplementary Figures for "Vacuolar pH regulates clathrin-mediated endocytosis through TORC1 signaling during yeast replicative aging"

### Figure S1

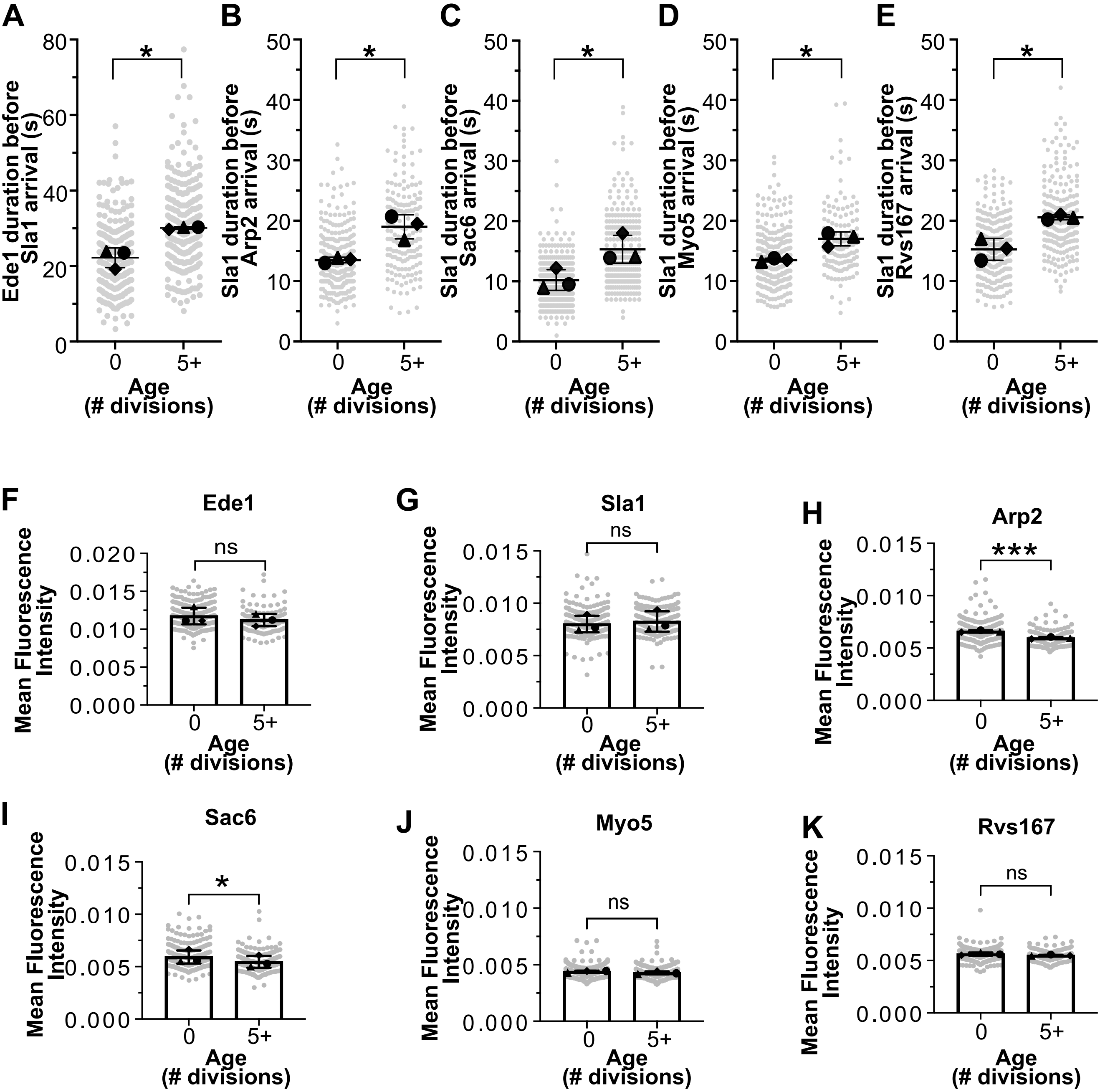

A
B
C
D
E
Ede1 duration before Sla1 arrival (s)
Sla1 duration before Arp2 arrival (s)
Sla1 duration before Sac6 arrival (s)
Sla1 duration before Myo5 arrival (s)
Sla1 duration before Rvs167 arrival (s)
*
Age (# divisions)
0
5+
F
Ede1
G
Sla1
H
Arp2
I
Sac6
J
Myo5
K
Rvs167
Mean Fluorescence Intensity
ns
***

### Figure S2

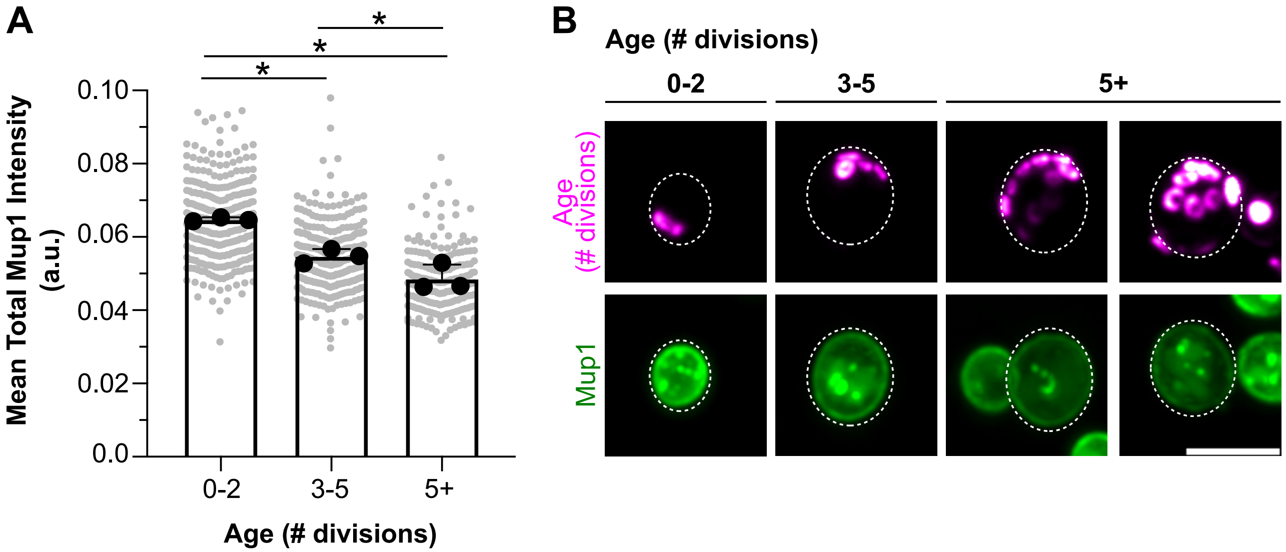
A
B
Mean Total Mup1 Intensity (a.u.)
0.10
0.08
0.06
0.04
0.02
0.0
0-2
3-5
5+
Age (# divisions)
*
*
*
Age (# divisions)
0-2
3-5
5+
Age (# divisions)
Mup1

### Figure S3

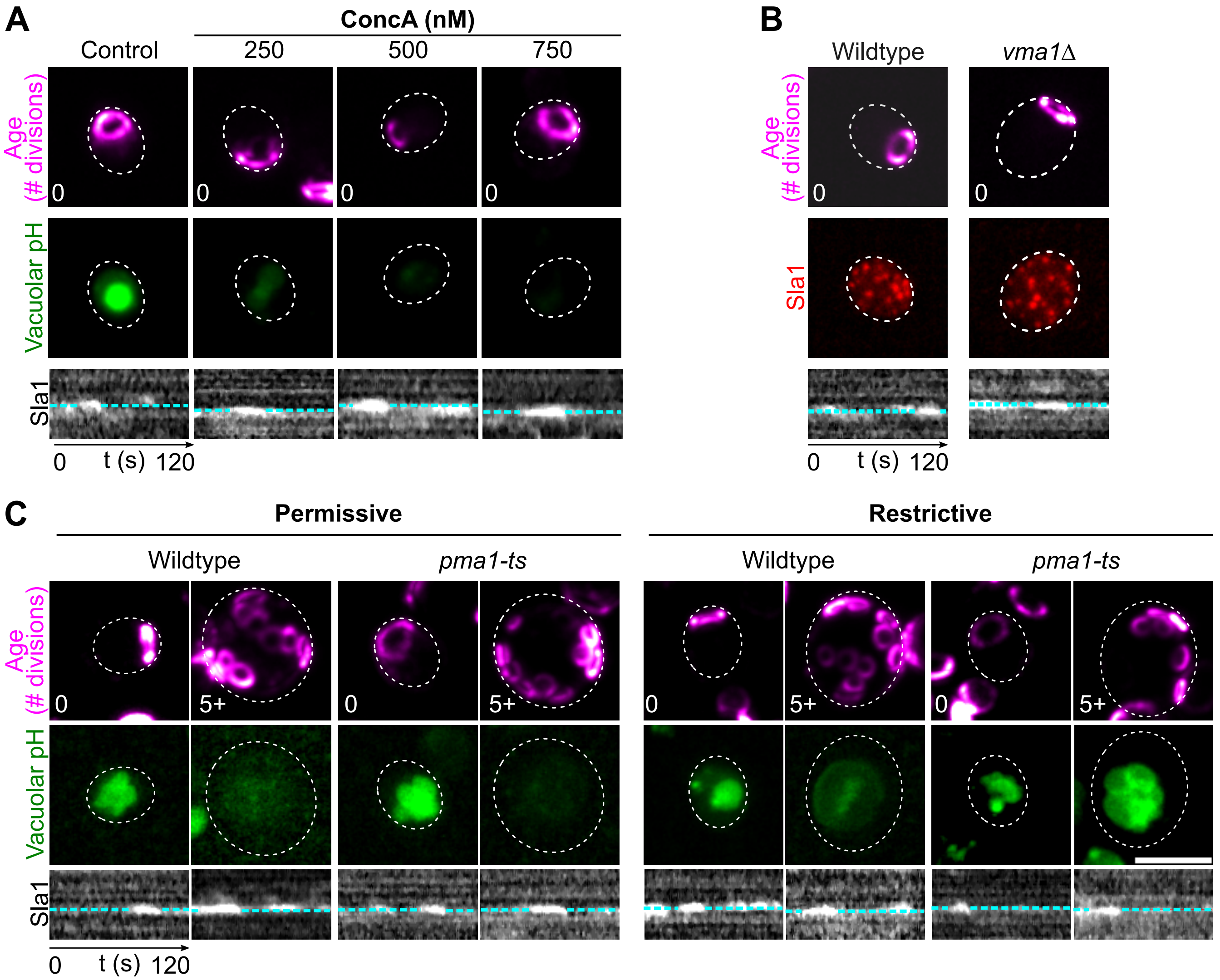

A
ConcA (nM)
Control
250
500
750
Age (# divisions)
0
0
0
0
Vacuolar pH
Sla1
0
t (s)
120
B
Wildtype
*vma1Δ*
Age (# divisions)
0
0
Sla1
Sla1
0
t (s)
120
C
Permissive
Restrictive
Wildtype
*pma1-ts*
Wildtype
*pma1-ts*
Age (# divisions)
0
5+
0
5+
0
5+
0
5+
Vacuolar pH
Sla1
0
t (s)
120

### Figure S4

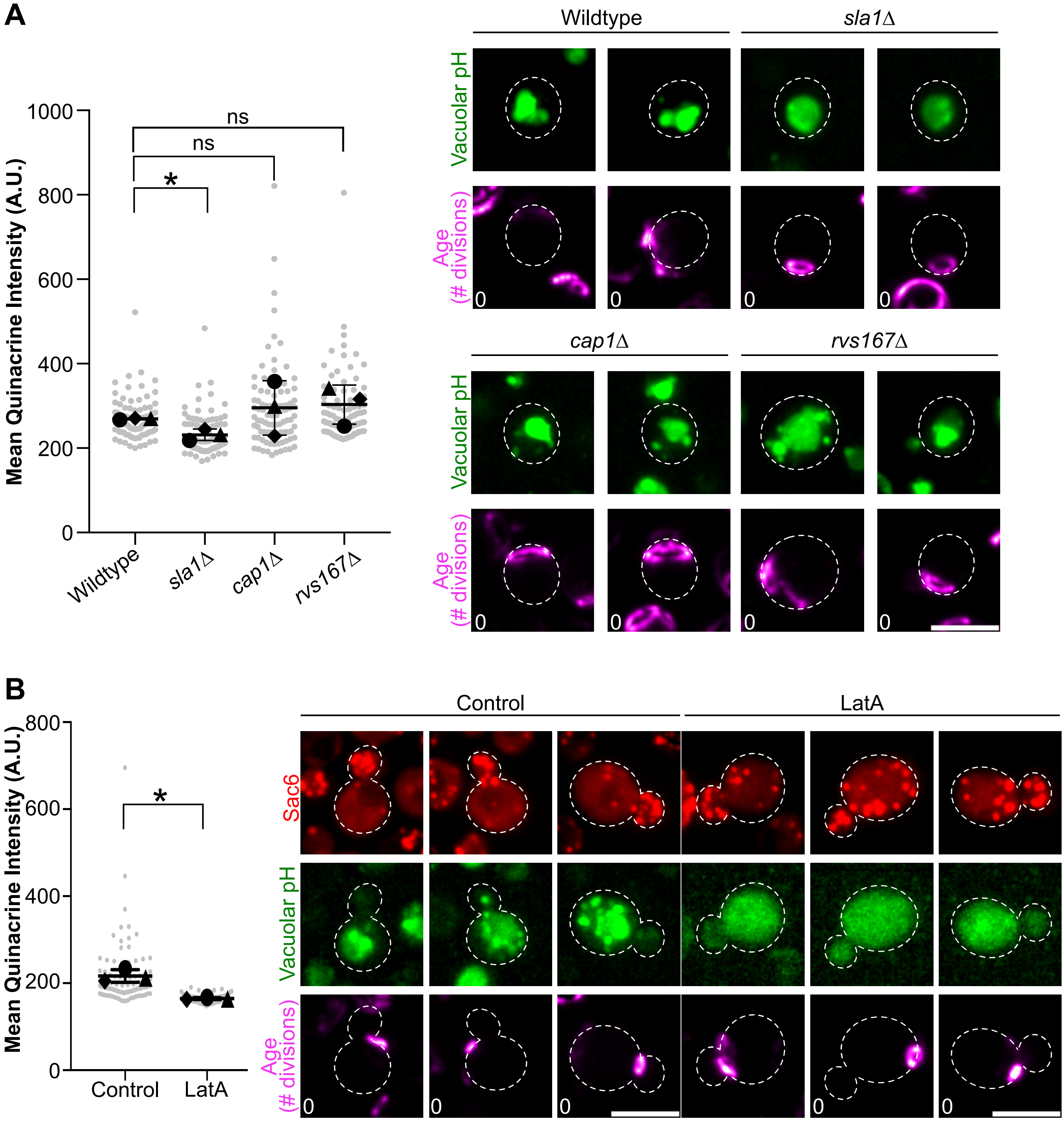
A
Mean Quinacrine Intensity (A.U.)
1000
800
600
400
200
0
ns
ns
*
Wildtype
sla1Δ
cap1Δ
rvs167Δ
Vacuolar pH
Age (# divisions)
0
B
Control
LatA
Sac6

Figure S5

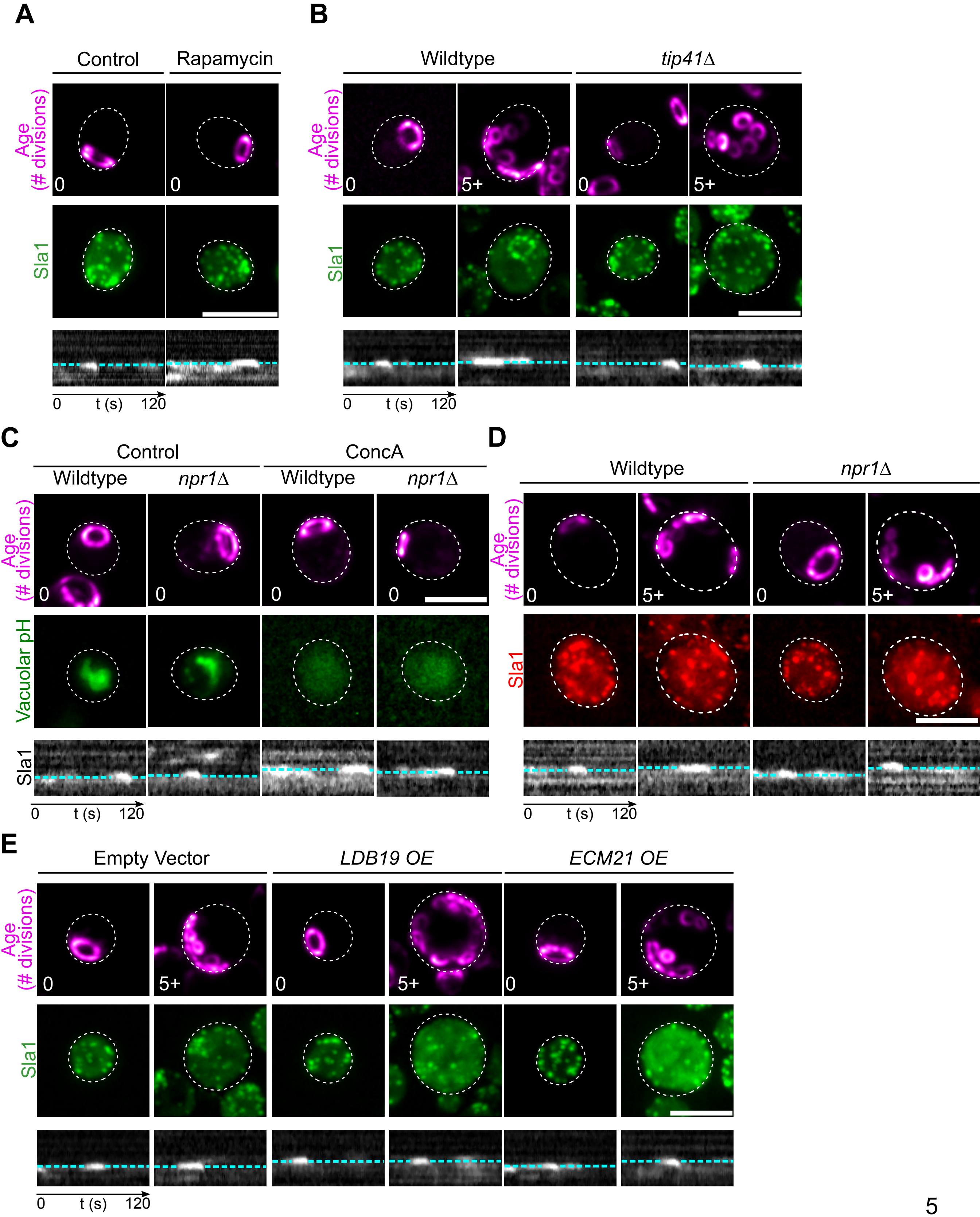
A
Control
Rapamycin
Age (# divisions)
0
Sla1
0
t (s)
120
B
Wildtype
tip41Δ
Age (# divisions)
5+
Sla1
C
Control
ConcA
Wildtype
npr1Δ
Vacuolar pH
D
E
Empty Vector
LDB19 OE
ECM21 OE
