## Supplemental Table 1 for "Vacuolar pH regulates clathrin-mediated endocytosis through TORC1 signaling during yeast replicative aging"

**Table S1. Yeast Strains, Plasmids and Oligonucleotides used in this study**

**A) Yeast Strains**

| ID | Genotype | Source | Construction Details |
|---|---|---|---|
| **BY4741** | *MATa his3Δ1 leu2Δ0 met15Δ0 ura3Δ0* | Brachmann et al, 1998 | NA |
| **SY2014** | *MATα ste3Δ306::LEU2 sst2Δ* | Anderson et al., 2003 | NA |
| **SY2625** | *MATa bar1Δ* | Anderson et al., 2003 | NA |
| **PMY178.1** | *MATα his3Δ1 ura3Δ0 leu2Δ lyp1Δ met15Δ0 can1Δ::STE2pr-spHIS5 EDE1-yomNeonGreen::NatMX SAC6-yomScarlet-I::cgLEU2 HTA2-TagBFP::HygMX* | Dr. D. Drubin | NA |
| **SLA1-GFP** | *MATa his3Δ1 leu2Δ0 met15Δ0 ura3Δ0 SLA1-GFP::spHIS5* | Yeast ORF-GFP collection (Huh et al., 2003.) | NA |
| **SAC6-GFP** | *MATa his3Δ1 leu2Δ0 met15Δ0 ura3Δ0 SAC6-GFP::spHIS5* | Yeast ORF-GFP collection (Huh et al., 2003) | NA |
| **MUP1-GFP** | *MATa his3Δ1 leu2Δ0 met15Δ0 ura3Δ0 MUP1-GFP::HIS3MX* | Yeast ORF-GFP collection (Huh et al., 2003) | NA |
| ***tip41Δ*** | *MATa his3Δ1 leu2Δ0 ura3Δ0 met15Δ0 tip41Δ::KANMX* | Yeast deletion collection (Giaever et al., 2002) | NA |
| ***pma1-ts*** | *MATα his3Δ1 leu2Δ0 ura3Δ0 met15Δ0 pma1-ts::NATMX* | Yeast Collection of Temperature-sensitive Strains (Li et al, 2011; Costanzo et al, 2016) | NA |
| ***vma1Δ*** | *MATα his3Δ1 leu2Δ0 ura3Δ0 met15Δ0 vma1Δ::NATMX* | Yeast deletion collection (Giaever et al., 2002) | NA |
| ***npr1Δ*** | *MATa his3Δ1 leu2Δ0 ura3Δ0 met15Δ0 npr1Δ::KANMX* | Yeast deletion collection (Giaever et al., 2002) | NA |
| ***oxr1Δ*** | *MATa his3Δ1 leu2Δ0 ura3Δ0 met15Δ0 oxr1Δ::KANMX* | Yeast deletion collection (Giaever et al., 2002) | NA |

| | | | |
|---|---|---|---|
| **MUY142** | *MATα can1Δ::STE2pr-Sp_his5 lyp1Δ his3Δ1 leu2Δ0 ura3Δ0 met15Δ0 VPH1-yEGFP::NATMX SLA1-tdTomato::URA3* | Mattiazzi Usaj et al., 2020 | NA |
| **MUY10** | *MATa his3Δ1 leu2Δ0 met15Δ0 ura3Δ0 SLA1-yomNeonGreen::NatMX* | This study | NA |
| **MUY110** | *MATα his3Δ1 ura3Δ0 leu2Δ0 lyp1Δ met15Δ0 can1Δ::STE2pr-spHIS5 HTA2-TagBFP::HygMX SLA1-yomNeonGreen::NatMX ARP2-yomScarlet-I::cgLEU2* | This study | *, pPM38 and ARP2mScarlet_F/R |
| **MUY111** | *MATα his3Δ1 ura3Δ0 leu2Δ0 lyp1Δ met15Δ0 can1Δ::STE2pr-spHIS5 HTA2-TagBFP::HygMX SLA1-yomNeonGreen::NatMX MYO5-yomScarlet-I::cgLEU2* | This study | *, pPM38 and MYO5mScarlet_F/R |
| **MUY112** | *MATα his3Δ1 ura3Δ0 leu2Δ0 lyp1Δ met15Δ0 can1Δ::STE2pr-spHIS5 HTA2-TagBFP::HygMX SLA1-yomNeonGreen::NatMX RVS167-yomScarlet-I::cgLEU2* | This study | *, pPM38 RVS167mScarlet_F/R |
| **MUY155** | *MATα his3Δ1 ura3Δ0 leu2Δ0 lyp1Δ met15Δ0 can1Δ::STE2pr-spHIS5 HTA2-TagBFP::HygMX SLA1-yomScarlet-I::cgLEU2* | This study | *, pPM38 and SLA1mScarlet_F/R |
| **MUY156** | *MATα his3Δ1 ura3Δ0 leu2Δ0 lyp1Δ met15Δ0 can1Δ::STE2pr-spHIS5 HTA2-TagBFP::HygMX SLA1-yomNeonGreen::NatMX EDE1-yomScarlet::cgLEU2* | This study | *, pPM38 and EDE1mScarlet_F/R |
| **MUY158** | *MATα his3Δ1 ura3Δ0 leu2Δ0 lyp1Δ met15Δ0 can1Δ::STE2pr-spHIS5 HTA2-TagBFP::HygMX SLA1-mCherry::caURA3* | This study | *, BA425v and SLA1_F/R |
| **MUY170** | *MATα his3Δ1 leu2Δ0 ura3Δ0 met15Δ vma1Δ::NATMX SLA1-mCherry::caURA3* | This study | *, BA425v and SLA1_F/R |
| **MUY182** | *MATa pma1-ts::NATMX his3Δ1 leu2Δ0 ura3Δ0 met15Δ0 SLA1-mCherry::caURA3* | This study | *, BA425v and SLA1_F/R |
| **MUY32** | *MATα his3Δ1 ura3Δ0 leu2Δ0 lyp1Δ met15Δ0 can1Δ::STE2pr-spHIS5 SAC6-yomScarlet-I::cgLEU2 HTA2-TagBFP::HygMX SLA1-yomNeonGreen::NatMX* | This study | **; MUY10 and PMY178.1 |
| **MUY64** | *MATα his3Δ1 ura3Δ0 leu2Δ0 lyp1Δ met15Δ0 can1Δ::STE2pr-spHIS5 SAC6-yomScarlet-I::cgLEU2 HTA2-TagBFP::HygMX SLA1-yomNeonGreen::NatMX tip41Δ::KanMX* | This study | **; MUY32 and *tip41*Δ |

| | | | |
|---|---|---|---|
| **MUY143** | *MATα his3Δ1 leu2Δ0 met15Δ0 ura3Δ0 MUP1-GFP::HIS3MX SLA1-tdTomato::URA3MX* | This study | **; MUP1-GFP and MUY142 |
| **MUY171** | *MATa his3Δ1 ura3Δ0 leu2Δ0 lyp1Δ met15Δ0 can1Δ::STE2pr-spHIS5 HTA2-TagBFP::HygMX npr1Δ::kanMX4 SLA1-mCherry::caURA3* | This study | **; MUY158 and *npr1*Δ |
| **MUY179** | *MATa his3Δ1 ura3Δ0 leu2Δ0 lyp1Δ met15Δ0 can1Δ::STE2pr-spHIS5 HTA2-TagBFP::HygMX oxr1Δ::kanMX4 SLA1-mCherry::caURA3* | This study | **; MUY158 and *oxr1*Δ |
| **MUY183** | *MATa, his3Δ1, ura3Δ0, leu2Δ0, lyp1Δ, met15Δ0, can1Δ::STE2pr-spHIS5, SLA1-yomNeonGreen::NatMX, HTA2-TagBFP::HygMX + p5819-KanMX-LEU2* | This study | Plasmid transformation |
| **MUY184** | *MATa, his3Δ1, ura3Δ0, leu2Δ0, lyp1Δ, met15Δ0, can1Δ::STE2pr-spHIS5, SLA1-yomNeonGreen::NatMX, HTA2-TagBFP::HygMX + pLDB19-KanMX-LEU2* | This study | Plasmid transformation |
| **MUY185** | *MATa, his3Δ1, ura3Δ0, leu2Δ0, lyp1Δ, met15Δ0, can1Δ::STE2pr-spHIS5, SLA1-yomNeonGreen::NatMX, HTA2-TagBFP::HygMX + pECM21-KanMX-LEU2* | This study | Plasmid transformation |

**Legend for Strain Construction Method: *, Homologous Recombination of PCR Product; ** Genetic Crossing; NA, not applicable**

**B) Plasmids**

| Plasmid | Description | Source/Reference |
|---|---|---|
| pPM45 | pKT-yomNeonGreen-NatMX | Dr. D. Drubin |
| pPM38 | pFA6a-yomScarlet-I-cgLEU2 | Dr. D. Drubin |
| BA425v | pkT-mCherry-CaURA3 | Dr. B. Andrews |
| p5819 | 2μ Empty Vector for MoBY-ORF 2.0 | MoBY-ORF Library 2.0 (Magtanong et al., 2011) |
| MoBY-ORF 2.0 *LDB19* | *2μ LDB19::KanMX* | MoBY-ORF Library 2.0 (Magtanong et al., 2011) |
| MoBY-ORF 2.0 *ECM21* | *2μ ECM21::KanMX* | MoBY-ORF Library 2.0 (Magtanong et al., 2011) |

**C) Oligonucleotides**

| ID | Sequence (5' to 3') |
|---|---|
| SLA1mScarlet_F | GCA AGC CAA CAT ATT CAA TGC TAC TGC ATC AAA TCC GTT TGG ATT CGC GGC CGC TCT AGA ACT AGT GG |
| SLA1mScarlet_R | GCC ATT TTC ACG AGT ATA AGC ACA GAT TGT ACG AAA CTA TTT CAT ATA GC GTT TAA ACG AGC TCG AAT TC |
| SLA1_F | CAAGCCAACATATTCAATGCTACTGCATCAAATCCGTTTGGATTCGGTGACGGTGCTGGTTTA |
| SLA1_R | TTGCCATTTTCACGAGTATAAGCACAGATTGTACGAAACTATTTCGATATCATCGATGAATTCG |
| RVS167mScarlet_F | CAG CAA GGT GTG TTT CCT GGG AAC TAC GTG CAA CTC AAC AAG AAC GCG GCC GCT CTA GAA CTA GTG G |
| RVS167mScarlet_R | CCG GTT TCC CAA CTG GGC TGT GTC ATC GAC AAG ACA GAG ATG GTG GAA TTC GAG CTC GTT TAA AC |
| MY05mScarlet_F | GAG TGA TGA CGA GGA GGC TAA CGA AGA TGA AGA GGA AGA TGA TTG GGC GGC CGC TCT AGA ACT AGT GG |
| MY05mScarlet_R | CGT TCA GGA AGA ACA ATG AGC GTA AAT GGT ACT AAT TAA AAT AAC GGC GGC GAA TTC GAG CTC GTT TAA AC |
| ARP2mScarlet_F | GGC AAG AAA GCG GGC CAT CTG CAA TGA CTA AAT TTG GTC CAA GA G CGG CCG CTC TAG AAC TAG TGG |
| ARP2mScarlet _R | CTG AGC GCC GTT AGA GGA TAG TTA TCC TTT TCT TGG TAT TTC AAG AAT TCG AGC TCG TTT AAA C |
| EDE1mScarlet _F | GCA ACT GGG ATC TAG AAG CCG CCA CTA ACT TTT TGT TGG ATA GTG CT G CGG CCG CTC TAG AAC TAG TGG |
| EDE1mScarlet _R | GCT CTA CCA CAT AAT CTT TGG GCG TAT TAC AAC ATG GGA AAC ACG ACG AAT TCG AGC TCG TTT AAA C |
| ARP2_UP | GCC GTT TTA GCT AGT ATC ATG GC |
| ARP2_DN | GGA ATG CCG TTA GAT ATG GC |
| EDE1_UP | GGG ATT TAC TGA AGA AGA AGC |
| EDE1_DN | CAA CTG GAC CAA ACC AAA ACC C |
| MYO5_UP | CTC TAA GGA AGG TTG GGT GC |

| | |
|---|---|
| MYO5_DN | GCT CGT CCG TTT TGC GAT GG |
| RVS167_UP | GGT GAT TTG TCT TTC CCT GCC G |
| RVS167_DN | CCT GCT TGA AGA TGT TGT GC |
| cgLEU2promoter_UP | TCC CGT ATA ATG TGT GCT CTG |
| cgLEU2promoter_DN | CAT GTG ACC TGT ATC GCT CA |
